# Alternative splicing in mechanically stretched podocytes as a model of glomerular hypertension

**DOI:** 10.1101/2024.07.17.603550

**Authors:** Francescapaola Mattias, Olga Tsoy, Elke Hammer, Alexander Gress, Stefan Simm, Chit Tong Lio, Sabine Ameling, Kerstin Amann, Leonie Dreher, Ulrich Wenzel, Tim Kacprowski, Markus List, Olga Kalinina, Karlhans Endlich, Jan Baumbach, Uwe Völker, Nicole Endlich, Felix Kliewe

**Author notes:** Corresponding author: Dr. Felix Kliewe, Institut für Anatomie und Zellbiologie, Universitätsmedizin Greifswald Friedrich-Loeffler-Str. 23c, 17489 Greifswald, Germany. These authors contributed equally (Nicole Endlich, Felix Kliewe).

## Abstract

**Background:** Alterations in pre-mRNA splicing play an important role in disease pathophysiology. However, the role of alternative splicing (AS) for podocytes in hypertensive nephropathy (HN) has not been investigated. The purpose of the Sys_CARE project was to identify AS events that play a role in the development and progression of HN.

**Methods:** Murine podocytes were exposed to mechanical stretch, after which proteins and mRNA were analyzed by proteomics, RNA-Seq and several bioinformatic AS tools.

**Results:** Based on transcriptomics and proteomics analysis we could observe significant changes in gene expression and abundance of proteins under mechanical stretch compared to unstretched conditions. By RNA-Seq, we identified over 3,000 alternative spliced genes after mechanical stretch, including all types of AS events. We found 17 genes that showed an AS event in four different splicing analysis tools. From these, we focused on *Myl6*, a component of the myosin protein complex, and *Shroom3*, an actin-binding protein crucial for podocyte function. We found two *Shroom3* isoforms that showed significant changes in expression upon mechanical stretch, which was verified by qRT-PCR and *in situ* hybridization. Furthermore, we observed an expression switch of two *Myl6* isoforms after mechanical stretch. This switch is accompanied by a change in a C-terminally located amino acid sequence.

**Conclusions:** In summary, mechanical stretch of cultured podocytes is an excellent model to simulate hypertensive nephropathy. In depth RNA-Seq analysis disclosed alternative splicing events, such as in *Shroom3* and *Myl6*, which may play a crucial role in the pathophysiology of hypertension-induced nephropathy.

## Introduction

The podocytes, highly differentiated visceral epithelial cells in the kidney glomerulus, play a crucial role in the maintenance of the glomerular filtration barrier. Hypertension is a contributing factor to the progression of chronic kidney disease (CKD) and the development of end-stage kidney disease.^1,2^ Hypertensive nephropathy (HN) is caused by high blood pressure and the instability of podocyte architecture, resulting from the effacement of podocyte foot processes (FPs) or podocyte detachment.^3^ As mature podocytes have limited ability to divide, damaged or lost podocytes cannot be replaced.^4,5^

In higher eukaryotes, alternative splicing (AS) is the main process for generating transcriptome and proteome diversity for tissue- and development-specific expression. It has been shown that pre-mRNAs from approximately 95% of human genes with multiple exons undergo AS, resulting in multiple protein isoforms with different structures and functions from a single gene.^6^

Several studies have shown that defects in AS regulation may be the initial trigger for various diseases, including cancer,^7–9^ cardiovascular diseases,^10^ neurodegenerative diseases,^11,12^ and neuromuscular diseases.^13^ Since the completion of the sequencing and annotation of the human genome, several next-generation sequencing techniques have become available. As a result, the number of diseases found to be associated with changes in alternative splicing has increased dramatically.

Previous studies have described individual AS events that affect the development of kidney injuries. For instance, intron retention and the presence of an early stop codon result in the production of soluble Klotho (sKlotho).^14,15^ Supplementation of sKlotho has demonstrated a nephroprotective effect in model animals with CKD.^16,17^ During the early stages of diabetic nephropathy (DN), the mRNA level of the VEGF-A_165_b splice isoform (vascular endothelial growth factor) increases to protect the glomerular filtration barrier.^18^ The podocyte-specific protein WT1 also has two characteristic splicing events: the inclusion or skipping of exon 5 and the alternative use of two splice donor sites between exons 9 and 10, leading to the expression of two major protein isoforms, WT1(+KTS) and WT1(-KTS).^19^ Mice lacking the WT1(-KTS) variant exhibit more severe kidney defects, while those lacking the WT1(+KTS) variant show disturbed podocyte function.^20^

However, no studies have yet been reported in the literature on the correlation between AS defects and HN. In this study, we used our *in vitro* hypertension mouse model^21^ to compare omics data, detect and quantify AS events, gene isoforms, and isoform switches in mechanically stretched podocytes. In this context, we identified several significantly alternatively spliced transcripts, and focused on Myl6 (Myosin light chain 6) and Shroom3 (Shroom family member 3). Myl6, a cellular motor protein that generates force for cellular movements, is highly expressed in podocytes. Genome-wide association studies have found that SNPs in SHROOM3 are associated with chronic kidney disease (CKD) and lower estimated Glomerular Filtration Rate (eGFR).^22–26^ Additionally, Shroom3 deficiency in mice leads to glomerular abnormalities, albuminuria, and changes in podocyte morphology, such as foot process effacement.^27,28^

In summary, we detected differential AS events that may contribute to the development of hypertension-related glomerulopathies. These findings identify potential biomarkers that could be useful in predicting the onset and progression of these diseases.

## Methods

Further methods can be found in the “Supplementary methods” section.

### Cell Culture

Conditionally immortalized podocytes (SVI; CLS Cell Line Service GmbH, Eppelheim, Germany) were handled as described previously.^21^

### Mechanical stretch experiments

Mechanical stretch experiments were performed according to our previous study.^21^ Differentiated mouse podocytes were seeded on flexible silicon membranes of a six well plate (Bioflex, Flexcell^®^ International Corporation, Burlington, NC, USA). The flexible silicone membranes were coated with collagen IV to facilitate cell attachment. After 3 days, the six well plate was mounted on a manifold connected to a custom-built stretch apparatus (NIPOKA GmbH, Greifswald, Germany), which induced cyclic variations in air pressure below and above atmospheric pressure. Cyclic pressure variations caused upward and downward motion of the silicone membranes. Pressure amplitude was chosen to give a maximum upward deflection of the membrane center of 6 mm (low stretch), being equal to an increase in membrane area by 11%, or 5% mean linear cell strain. For the high stretch experiments, experiments were performed with a higher amplitude (8 mm). Cycle frequency was adjusted to 1 Hz.

### Ang II-induced hypertensive mice

Induction of Ang II-induced hypertension has been described previously.^29,30^

### Liquid chromatography-mass spectrometry (LC-MS)

Mechanically low and high stretched podocytes of three independent bioreplicates were harvested in Trizol and treated according to manufacturer’s protocol. Protein was obtained from the corresponding phase by isopropanol precipitation. Protein pellet was reconstituted in Tris-HCl (10 mM, pH 7.5) with 5% SDS. Protein concentration was determined by BCA assay (Pierce/Thermo Fisher Scientific). We used an adapted bead based SP3 protocol^31^ for removal of contaminants and proteolytic digestion. Briefly, five µg protein were reduced by 2.5 mM dithiothreitol for 30 min at 37°C and alkylated with 10 mM iodoacetic acid for 30 min at 37°C in the dark before digestion by trypsin (Promega, Walldorf, Germany) in a protein to enzyme ratio 25:1 overnight at 37°C. Peptides were separated by LC (Ultimate 3000, Thermo Electron, Bremen, Germany) before data-independent acquisition of MS data on an Exploris 480 mass spectrometer (Thermo Electron). MS data were analyzed via the DirectDIA algorithm implemented in Spectronaut (v17, Biognosys, Zurich, Switzerland) using an Uniprot database (version 2021_2) limited to *Mus musculus* (n=17063). Carbamidomethylation at cysteine was set as static modification, oxidation at methionine and protein N-terminal acetylation were defined as variable modifications, and up to two missed cleavages were allowed. Proteins were only considered for further analyses, if two or more unique+razor peptides were identified and quantified per protein. Further data analysis was performed as reported earlier.^32^ Detailed description of data acquisition and search parameters are provided in Supplemental Table S2.

### RNA-seq and AS event analysis

RNA-Seq raw reads for low-stretch, high-stretch, and unstretched podocytes were mapped to the mouse reference genome mm39 with the genome annotation from Ensembl version 104 using STAR version 2.7.5c.^33^ We used the default parameters with a 2-pass mode recommended for alternative splicing analysis. Gene read counts were obtained by the featureCounts tool included in the Subread 2.0.0.^34^ Read counts were further normalized and used for differential gene expression by DESeq2 version 1.38.3.^35^ For isoform switch analysis, we prepared transcript counts using salmon version 1.9.0^36^ with the transcript annotation based on Ensembl version 104 and ran the R package IsoformSwitchAnalyzeR version 1.20.0^37^ with the DESeq2 algorithm. For alternative splicing analysis, we used leafcutter version 0.2.9,^38^ rMATS version 4.1.2,^39^ and Whippet version 0.11.1 (with julia 1.6.6)^40^ with the default parameters. We defined significantly differentially used events as events with a p-value < 0.05 (for leafcutter, rMATS, and Whippet).

### Statistical and bioinformatic analysis

Data were analyzed by Gene Set Enrichment Analysis (GSEA). For the analysis of the differentially abundant proteins and transcripts R scripts including the libraries <u>EnhancedVolcano</u> and <u>ComplexHeatmap</u> were used for the visualization. The heatmaps are used to represent the abundance of proteins as a ratio from treatment versus control of specific biological pathways from the GeneOntology (GO). To detect the enriched or overrepresented pathways and their protein protein interactions the dotplots from the GO biological processes were created using the library clusterProfiler.^41^

The GraphPad Prism 9 software was used for statistical analysis of experimental data and preparation of graphs. Scatter plots indicate individual units used for statistical testing (samples, cells or replicates), as specified in respective figure legends. Data are given as means ±SD or ±SEM, analyzed by unpaired *t* test with repeated measurements (n). For multiple groups statistical analyses were done by ANOVA with a Benjamini-Hochberg post-hoc test. Statistical significance was defined as p < 0.05 and significance levels are indicated as * p < 0.05, ** p < 0.01, *** p < 0.001, **** p < 0.0001 and non-significant (ns) in respective figure panels. The number of independent experiments and analyzed units are stated in the figure legends.

## Results

### Proteomics and transcriptomics of mechanically stretched podocytes

The objective of this study was to determine whether mechanical stretch regulates alternative splicing. To this end, podocytes were cultured on flexible silicone membranes and stretched at low or high stretch conditions for three days, which resulted in a strong reorganization of the F-actin (Fig. 1A-B). The mRNA and proteins were isolated and analyzed using complementary transcriptome (RNA-Seq) and proteome analysis (LC-MS/MS) (Fig. 1C).

**Fig. 1:**
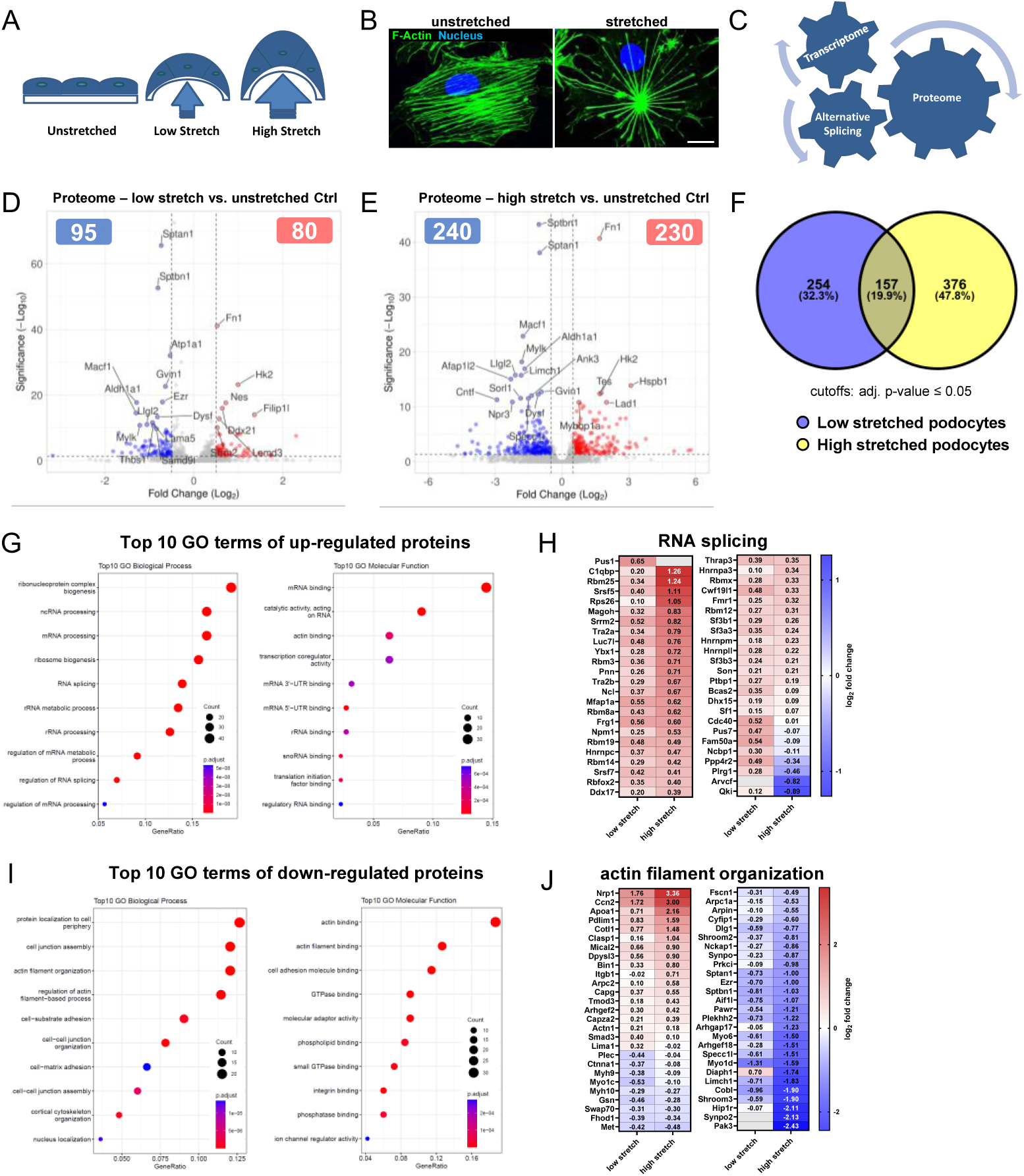
Proteomics of mechanically stretched podocytes. (A) Schematic representation of the different stretch conditions. (B) F-actin staining (shown in green) of unstretched and mechanically stretched podocytes. Nuclei are shown in blue. Scale bar represents 25 µm. (C) RNA and proteins from the stretched podocytes were analyzed by RNA-Seq and LC-MS/MS to investigate the mutual influence of the transcriptome and alternative splicing on the proteome. (D, E) Volcano plot displaying the significantly changed proteins upon (D) low and (E) high stretch conditions. Given are proteins with a log2 fold-change ≥ 0.5 and adjusted p-value ≤ 0.05. The 20 most significantly regulated proteins after mechanical stretch are labeled. (F) Venn diagram depicts overlap of 157 proteins that are significantly regulated at low and high stretch conditions (cutoffs: adj. p-value ≤ 0.05). (G) Top 10 GO (Gene Ontology) clusters in podocytes, based on significantly up-regulated proteins in mechanically stretched podocytes compared to unstretched Ctrl. (H) Heatmap (LC-MS/MS based relative protein level) of regulated “RNA splicing” proteins (GO term) in low and high stretched podocytes. Data are represented as log2 fold change, a p-value < 0.05 was considered as significant. Red: up-regulated; blue: down-regulated compared to controls. (I) Top 10 GO (Gene Ontology) clusters in podocytes, based on significantly down-regulated proteins in mechanically stretched podocytes compared to unstretched Ctrl. (J) Heatmap (LC-MS/MS based relative protein level) of regulated “actin filament organization” proteins (GO term) in low and high stretched podocytes. Data are presented as in Fig. H.

Proteome analysis revealed significant alterations upon mechanical stretch. A total of 175 proteins showed different levels under low stretch conditions (80 increased and 95 decreased). Under high stretch conditions, 470 proteins were altered (230 increased and 240 decreased) compared to the unstretched condition (cutoffs: adj. p-value < 0.05 and log_2_ fold change ≥ 0.5) (Fig. 1D-E; Supplementary Data S1). These results show that higher mechanical stress leads to a strong regulation of proteins in podocytes. From 787 differentially abundant proteins (cutoff: adj. p-value < 0.05), 157 (20%) are found in both low and high stretch conditions (Fig. 1F).

Gene set enrichment analysis (GSEA) for gene ontology (GO) terms revealed an up-regulation of proteins associated with mRNA processing and RNA splicing in mechanically stretched podocytes in comparison to unstretched podocytes (Fig. 1G-H, Fig. S1A, Fig. S1C-D). In contrast, the majority of proteins with decreased intensity upon stretch were cytoskeleton and actin-binding proteins (Fig. 1I-J, Fig. S1B, Fig. S1E-F).

Transcriptome analyses using RNA sequencing confirmed that increased mechanical stretch results in a higher number of differentially expressed genes. We observed 20% more down- and 106% more significantly up-regulated genes under high stretch conditions, compared to low stretch conditions (Fig. 2A-B). In total, we identified more than 1300 significantly regulated genes after mechanical stretch compared to unstretched podocytes (Fig. 2C; Supplementary Data S2). The GSEA of the transcriptome showed a similar enrichment of GO-Terms compared to the proteome (Fig. S2). Notably, several of the regulated targets are long non-coding RNAs (lncRNAs), that might impact gene regulation (Fig. 2C; Supplementary Data S2).

**Fig. 2:**
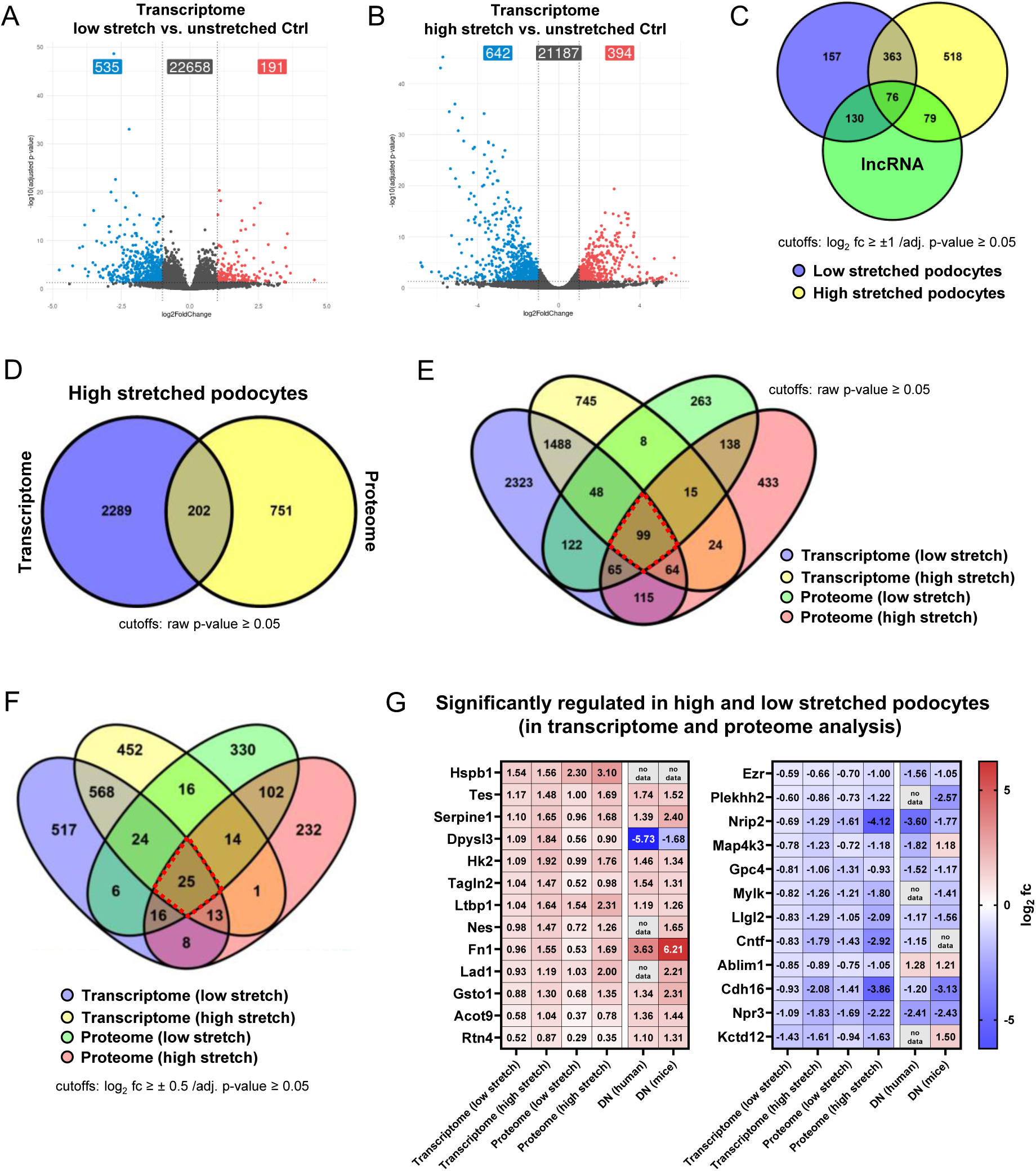
RNA-Seq data of mechanically stretched podocytes. (A, B) Volcano plot displaying the significantly changed genes at (A) low and (B) high stretch conditions. Given are genes with a log2 fold-change ≥ 1 and adjusted p-value ≤ 0.05. (C) Venn diagram shows overlap of 76 proteins that are significantly regulated at low and high stretch conditions and belong to the lncRNAs (cutoffs: adj. p-value ≤ 0.05 and log2 fold change ≥ 1). (D) Venn diagram of significantly regulated genes/proteins in high stretched podocytes (cutoffs: raw p-value ≤ 0.05). (E) Venn diagram depicts overlap of 99 proteins that are significantly regulated at low and high stretch conditions in the transcriptome analysis as well as in the proteome (cutoffs: raw p-value ≤ 0.05). (F) Venn diagram with higher thresholds depicts overlap of 25 proteins that are significantly regulated at low and high stretch conditions in the transcriptome analysis (cutoffs: adj. p-value ≤ 0.05 and log2 fold change ≥ 0.5) as well as in the proteome (cutoffs: adj. p-value ≤ 0.05). (G) Heatmap of significantly regulated genes/proteins in low and high stretched podocytes. In addition, the expression of the candidates in DN patients or DN mice is shown (data from Nephroseq). Data are represented as log2 fold change, a p-value < 0.05 was considered as significant. Red: up-regulated; blue: down-regulated compared to controls.

We further investigated which proteins are differentially regulated at the RNA as well as at the protein level (Fig. 2D-E). Our analysis identified 25 targets that were significantly regulated under both low and high stretch conditions on transcript as well as on protein level (Fig. 2F-G). Changes in the abundance of proteins and RNA were significantly higher at high stretch conditions in comparison to low stretch conditions. For example, the extracellular matrix protein fibronectin (*Fn1*), which is essential for podocyte adhesion,^42^ was found to be expressed 1.6-fold (at the RNA level) and even 3.2-fold (at the protein level) higher under high stretch conditions than under low stretch conditions (Fig. 2G). Furthermore, other known podocyte injury markers, such as Serpine1 or ezrin, were also significantly regulated at the mRNA and protein level, confirming mechanical stretch as a model for hypertension.

The regulation of the expression of various candidates was verified by qRT-PCR (Fig. S3). Interestingly, most of the genes/proteins that were regulated in our *in vitro* hypertension model were also significantly regulated *in vivo* in patients with diabetic nephropathy (DN), which is often associated with glomerular hypertension, or in DN mice (Fig. 2G).

### Alternative splicing in mechanically stretched podocytes

To identify alternative splicing (AS) events and decrease the number of false positive predictions, we used several AS detection tools (LeafCutter,^38^ rMATS,^39^ and Whippet.^40^ Additionally, we analyzed isoform switches using IsoformSwitchAnalyzeR^37^ (Fig. 3A). We found a wide variety of splicing events. The most frequent event was exon skipping (between 63-82% of all AS events), followed by intron retention and alternative 5′ or 3′ splice sites (Fig. 3B). We detected 290 alternative spliced genes detected using three different splice analysis tools (Fig. 3C). Of these 290 genes, 17 candidates also showed a significant isoform switch, including *Abca3*, *Arhgap22*, *Cenpi*, *Ikbkg*, *Myl6*, *Os9*, *Pja2*, *Prrc2b*, *Rgs12*, *Rps24*, *Rusc2*, *Shroom3*, *Slc27a1*, *Snx14*, *Sptan1*, *Tnnt2* and *Zfp219* (Fig. 3C-F, Fig. S4). To prioritize, we examined the renal and glomerular expression of these genes (Fig. 3D-E; Fig. S5). Furthermore, we screened which candidates were also present in the proteome, suggesting a podocyte-specific expression, and which showed a glomerular upregulation *in vivo* under hypertensive conditions (Fig. 3F). Based on these criteria, we focused on Myl6 and Shroom3.

**Fig. 3:**
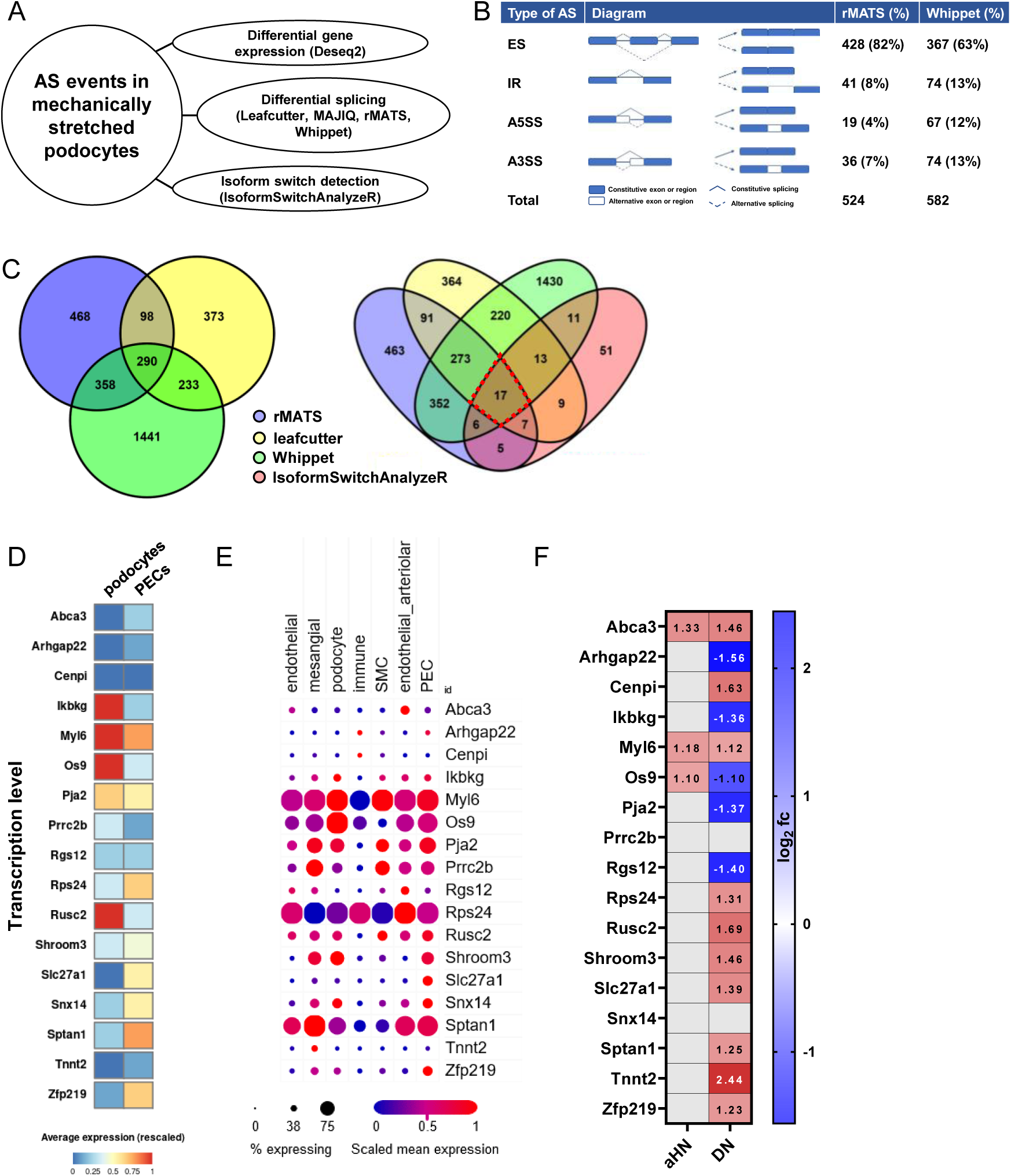
Alternative spliced transcripts in mechanically stretched podocytes. (A) Procedure of the AS event analysis. (B) Overview of the found AS types: ES (exon skipping); IR (intron retention); A5SS (Alternative 5′ splice site); A3SS (Alternative 3′ splice site). Only those AS events were considered which were significantly and have a deltaPSI > 0.1 (C) Venn diagram depicts overlap of 290 genes that showed a significant splice event in rMATS, leafcutter and Whippet. When filtering more stringently for the transcripts that also show an isoform switch (by using IsoformSwitchAnalyzer), 17 candidates remain. (D) Database analysis using the Kidney Cell Explorer by Ransick et al.^77^ based on a single-cell RNA sequencing data set of murine kidneys to determine podocyte-specific expression. Figure S6 shows expression pattern from all kidney cell fractions. Color code: Red means a high average expression; Blue: low expression. (E) Single-cell RNA-Seq data of mouse glomerulus (published from Chung et al.^78^ and available at https://singlecell.broadinstitute.org/single_cell). (F) mRNA expression level of renal glomeruli from mice with arterial hypertension (aHN) or human patients with diabetic nephropathy (DN). Data were taken from Nephroseq Research Edition (Ann Arbor, University of Michigan) and are represented as log2 fold change, a q-value < 0.05 was considered as significant. Red: up-regulated; blue: down-regulated compared to control.

### *Myl6* is alternative spliced in mechanically stretched podocytes

Myosin light chain 6 (Myl6), a component of the myosin motor protein complex, is highly expressed in cultured *in vitro* and *in vivo* podocytes (Fig. 4A-B and Fig. S6A). RNA-Seq data confirmed that *Myl6* is the most expressed myosin in podocytes (43 % of all myosins), which shows the importance of MYL6 for podocytes. (Fig. 4C). However, the *Myl6* expression is slightly, but not significantly increased after mechanical stretch (Fig. 4D). Furthermore, we have not observed a significant change of the MYL6 level in patients with hypertensive or diabetic nephropathy (Fig. 4E).

**Fig. 4:**
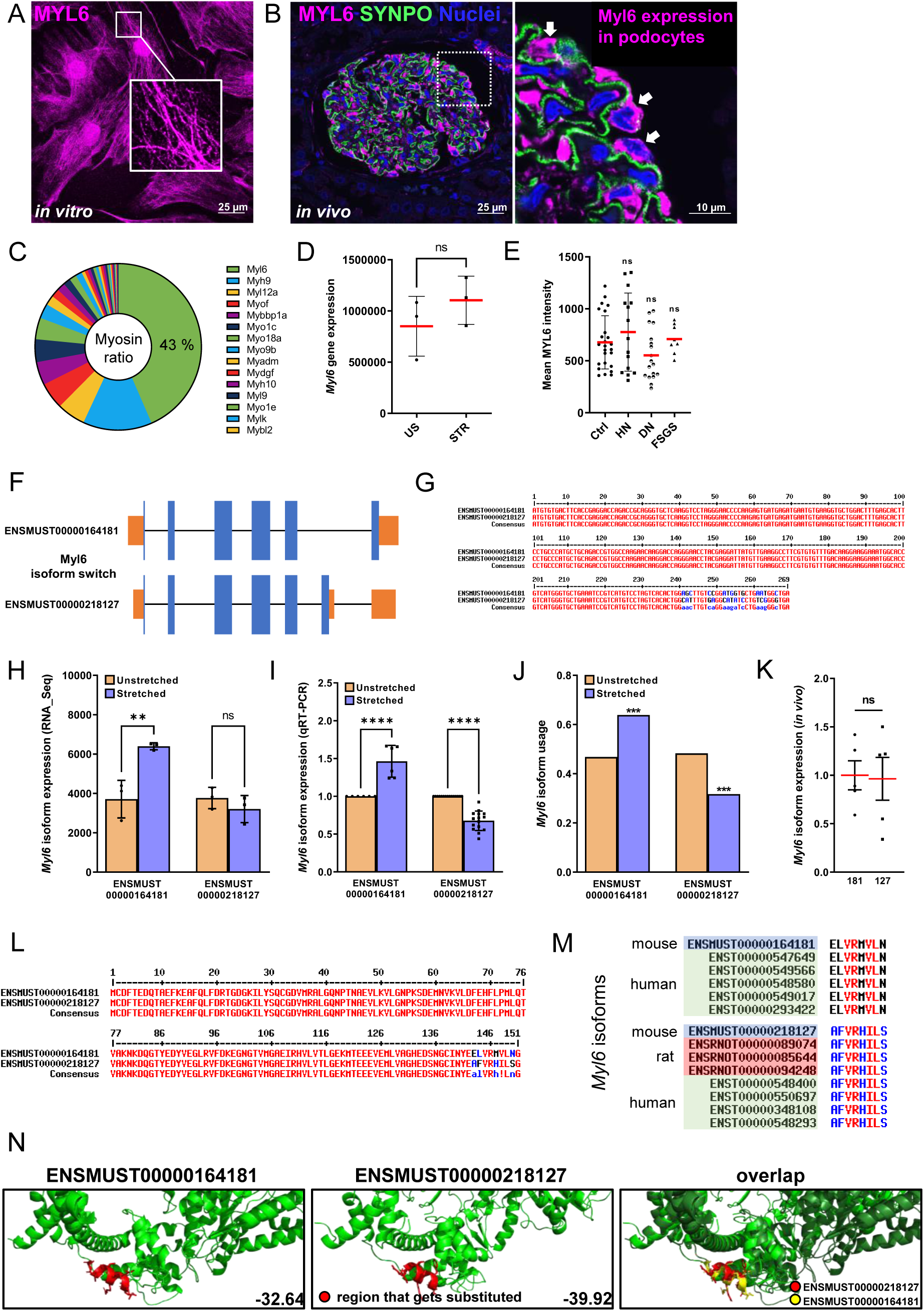
***Myl6* coded transcripts are alternative spliced in mechanically stretched podocytes.** (A) Immunofluorescence staining of MYL6 (magenta) showed high abundance in murine cultured podocytes. Scale bar represents 25 μm. (B) MYL6 (shown in magenta) is also expressed *in vivo* from podocytes (marked by white arrow heads). Synaptopodin is shown in green. Nuclei were stained with Hoechst (blue). Scale bars represent 25 µm and 10 µm (magnification) respectively. (C) Overview of the expression distribution of all myosin proteins found in podocytes. (D) *Myl6* gene expression in unstretched (US) und stretched (STR) podocytes. (E) Mean MYL6 fluorescence intensity in Ctrl, HN, DN and FSGS patients. Each dot represents the mean from one individual glomerulus (n≥15). (F) Schematic overview of the two Myl6 isoforms found in murine cultured podocytes. Blue: coding exons; Orange: untranslated regions. (G) Sequence alignment of coding exons of both *Myl6* isoforms. (H) *Myl6* isoform expression based on RNA-Seq data. (I) Validation of *Myl6* isoform expression by qRT-PCR. (J) *Myl6* isoform usage based on IsoformSwitchAnalyzer data. (K) *In vivo Myl6* transcript expression in isolated mouse glomeruli determined by qRT-PCR. (L) Protein sequence alignment of both MYL6 isoforms. (M) Amino acid switch from ELVRMVLN to AFVRHILS is well conserved in mouse, rat, and human. (N) Protein structural analyses revealed that amino acid substitution weakens the actin interaction surface. Data are presented as means ± SD. ** *p*<0.01; *** *p*<0.001; **** *p*<0.0001; ns, not significant.

We further studied the expression of isoforms of *Myl6*. By RNA sequencing and mass spectrometry we were able to detect two isoforms (Fig. 4F and Fig. S6B), which were differentially expressed in stretched versus unstretched conditions (Fig. 4H-J). However, the basic expression of both isoforms is almost identical *in vitro* under unstretched conditions (Fig. 4I) and *in vivo* (Fig. 4K). The two isoforms have the same length but differ in their C-termini (Fig. 4F-G). Isoform 2 (ENSMUST00000218127/NM_001317218) contains an alternate exon in the 3’ coding region, resulting in a frameshift and a novel 3’ UTR. This results in an amino acid switch from <u>EL</u>VR<u>MVLN</u> to <u>AF</u>VR<u>HILS</u> (Fig. 4L). These motifs are well conserved in mice, rats, and humans (Fig. 4M). Using the protein structure analysis tool StructMAn^43^ on these Myl6 isoforms, it was found that this amino acid substitution affects the actin binding domain (ABD) and weakens the actin interaction surface (Fig. 4N). Therefore, the isoform that increases after mechanical stretch has a higher potential for interaction with actin, which could benefit in withstanding increased mechanical stretch.

### Alternative splicing of *Shroom3 in vitro* and *in vivo*

Four Shroom3 isoforms were identified in cultured murine podocytes by RNA-Seq (Fig. 5A), with varying expression levels (Fig. 5C). Surprisingly, mouse podocytes express protein isoform 1 (encoded by ENSMUST00000113055/ NM_015756) at a very low level. This isoform is the longest and the only transcript that includes the PDZ domain and in human the intron with a GFR-associated SNP (Fig. 5A-B, Fig. S7). The transcript ENSMUST00000113054 (NM_001077595) encodes protein isoform 2, which has the same protein sequence like transcript variant ENSMUST00000113051 (NM_001077596), but differs in the 5’ UTR (Fig. 5A-B, S7). The isoform ENSMUST00000113054 was expressed at the highest level of all isoforms and showed a significant decrease in expression after mechanical stretch (Fig. 5C, E). Isoform ENSMUST00000225438 (XM_006534966) was up-regulated after mechanical stretch (Fig. 5C-E). Downstream analysis by IsoformSwitchAnalyzeR showed for isoforms ENSMUST00000113054 and ENSMUST00000225438 a significantly changed usage after mechanical stretch (Fig. 5D), which was confirmed by qRT-PCR (Fig. 5E). By qRT-PCR of murine glomeruli, we could also impressively show that the *Shroom3* isoform ENSMUST00000113054 is the dominant renal glomerular *Shroom3* isoform *in vivo* (16-fold higher expressed than ENSMUST00000225438) (Fig. 5F). This was confirmed by *in situ* hybridization experiments (Fig. 5I and Fig. S8). Additionally, *Shroom3* isoform ENSMUST00000113054 is exclusively expressed in the renal glomerulus (Fig. 5G). This finding was quantified with qRT-PCR, which showed that *Shroom3* ENSMUST00000113054 was around 12-fold enriched in isolated glomeruli in comparison with total kidney fractions (Fig. 5G). In a published isolated podocyte versus bulk glomerulus mass spectrometry dataset, podocyte-specificity of SHROOM3 in comparison to whole glomeruli could be verified (Fig. 5H).

**Fig. 5:**
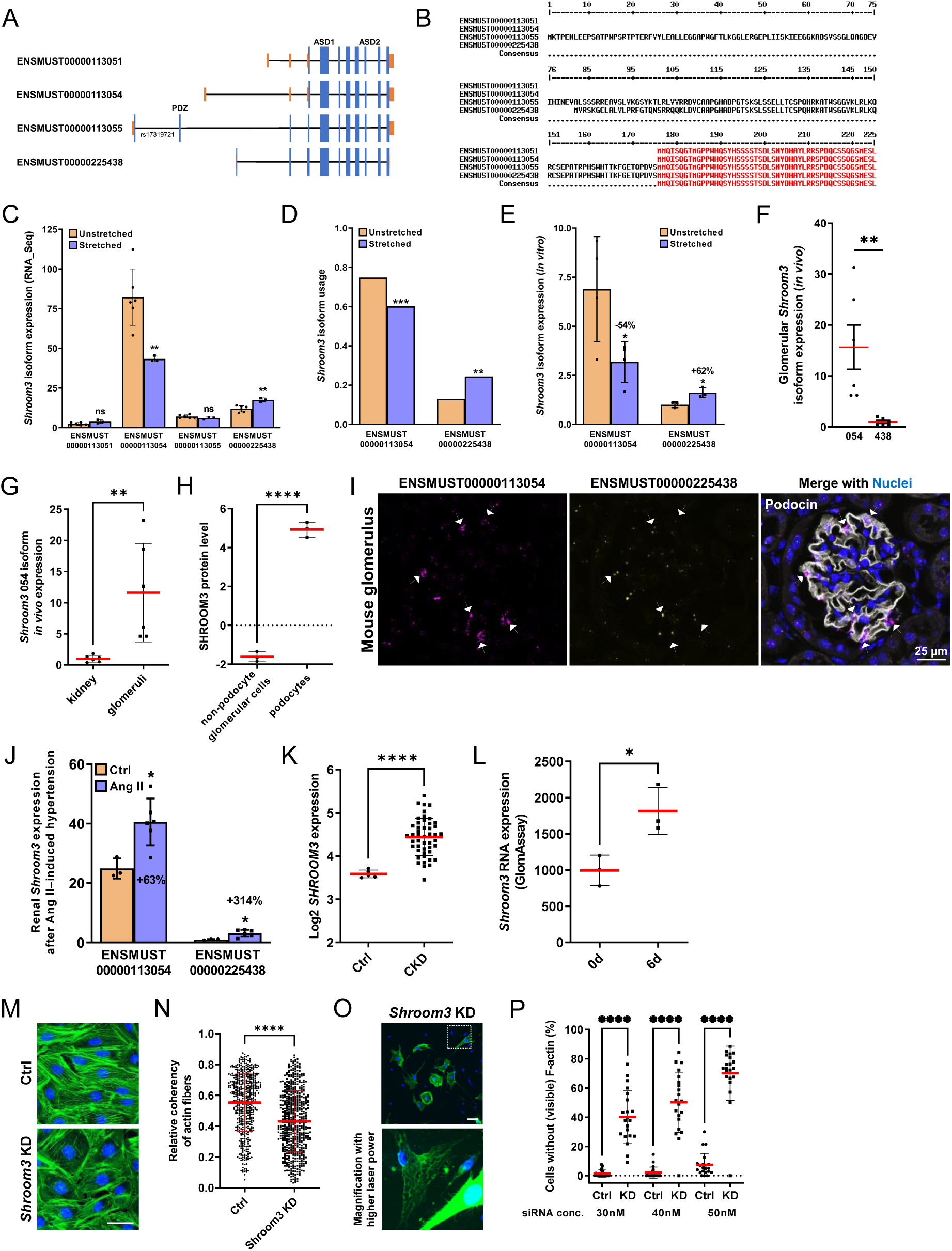
Alternative spliced isoforms of *Shroom3* in mechanically stretched podocytes. (A) Schematic overview of all *Shroom3* isoforms found in murine cultured podocytes. PDZ domain and ABD (actin binding domain) are marked. Blue: coding exons; Orange: untranslated regions (UTRs). (B) Protein sequence alignment of all SHROOM3 isoforms. Shown are the first 225 amino acids; complete sequence is shown in Fig. S7. (C) *Shroom3* isoform expression based on RNA-Seq data. (D) *Shroom3* coded isoform usage based on IsoformSwitchAnalyzer data. (E) Validation of *Shroom3* isoform switch by qRT-PCR. (F) qRT-PCR confirmed that transcript ENSMUST00000113054 is significantly higher expressed in *in vivo* glomeruli than ENSMUST00000225438. (G) *Shroom3* isoform 054 expression in whole kidney and isolated glomeruli. qRT-PCR experiments were normalized to the kidney samples and *Gapdh*. (H) Podocyte proteome data showed an enrichment of SHROOM3 protein level in podocytes compared to non-podocyte glomerular cells. Data from Rinschen et al.^79^. (I) Both isoforms of *Shroom3* were detected at different levels by *in situ* hybridization. Immunostaining of podocin is shown in white. Nuclei stained with Hoechst (blue). Scale bar represents 25 μm. (J) mRNA expression of *Shroom3* in murine kidneys with angiotensin II-induced hypertension (n=6) compared to Ctrl (n=3). (K) Analysis of human glomerular disease data^80^ (Nephroseq database) further substantiated increased expression on mRNA levels of *SHROOM3* in CKD patients (n = 47) compared to healthy controls (Ctrl) (n = 5). (L) *Shroom3* mRNA level of isolated glomeruli until 6 days of cultivation and transdifferentiation. The mRNA fold enrichment was normalized to day 0 and showed a significant increase of *Shroom3* expression. (M) Immunofluorescence staining of F-actin (green) of *Shroom3* KD and control (Ctrl) podocytes (transfected with non targeting control siRNA). Knockdown of *Shroom3* leads to a formation of an unorganized actin cytoskeleton. Nuclei were stained with DAPI (blue). Scale bar represents 100 μm. (N) The relative coherency of actin fibers was measured using OrientationJ (n = 3; each 20 analyzed cells; means ± SD). (O) *Shroom3* KD leads to dramatic reduction in F-actin formation. F-Actin is shown in green. Nuclei were stained with DAPI (blue). Scale bar represents 100 μm. (P) Percentage of cells without (visible) F-actin compared to controls (transfected with negative scrambled siRNA). * *p*<0.05; ** *p*<0.01; *** *p*<0.001; **** *p*<0.0001;ns, not significant.

To find out whether *Shroom3* is also regulated in mice suffering from hypertension, we analyzed the mRNA levels of *Shroom3* in murine kidneys with angiotensin II-induced hypertension (Fig. 5J). We observed that hypertensive mice showed a significant increase of *Shroom3* mRNA compared to the control mice (Fig. 5J). Furthermore, in human CKD patients and in our dedifferentiation GlomAssay, we observed a significant up-regulation of SHROOM3 (Fig. 5K and L).

To study the importance of *Shroom3* for podocytes, *Shroom3* was knocked down in cultured mouse podocytes using specific siRNA. Knockdown efficiency was validated by qRT-PCR (Fig. S9A). After treatment with specific siRNA against *Shroom3*, we observed a reorganization of the actin fibers specifically in *Shroom3* knockdown podocytes (*Shroom3* KD) (Fig. 5M). In contrast, we found mostly transversal stress fibers in the control cells as measured with the OrientationJ plugin^44^ for ImageJ (Fig. 5M). The relative coherence of the analyzed actin fibers was significantly reduced by 28% in *Shroom3* KD podocytes compared to the control (Fig. 5N). Interestingly, the total mRNA amount of actin (*Actb*) was significantly higher in *Shroom3* KD podocytes compared to the Ctrl (Fig. S9B). In contrast, the expression of intermediate filament and microtubule markers was not affected by *Shroom3* KD (Fig. S9B). Furthermore, we observed that with increasing siRNA concentration and the associated stronger knockdown effect, there is a dramatic reduction in F-actin formation (Fig. 5O-P and Fig. S9D). The proportion of cells without (visible) F-actin increased to over 70% at higher siRNA concentration. In contrast, the control cells transfected with the same concentration of negative scrambled siRNA do not show a higher proportion of cells without visible F-actin (Fig. 5P and Fig. S9D). The loss of SHROOM3 appears to result in an impaired polymerization of G-actin monomers to F-actin.

## Discussion

Podocyte damage and detachment caused by mechanical stress is a common issue in hypertension-induced glomerular disease.^21,45^ One of the cellular mechanisms that allow podocytes to adapt to these conditions is alternative splicing, a process that enables a single gene to produce multiple protein isoforms with distinct functions.^46,47^ This process is particularly important in podocytes, as they must dynamically reorganize their cytoskeleton, which is essential for maintaining podocyte structure and function, in response to mechanical stress, such as increased glomerular pressure in hypertension.

Our initial findings demonstrated that our *in vitro* hypertension model accurately reflects podocyte stress conditions. The mechanically stretched podocytes exhibited significant regulation of numerous known podocyte injury markers, including fibronectin, serpine1, and ezrin.^42,48–51^ These proteins serve as important markers of podocyte stress, reflecting the cellular response to mechanical stretch, such as increased glomerular pressure due to hypertension. Their regulation is associated with processes such as fibrosis or extracellular matrix and cytoskeleton remodeling, which are key features of chronic kidney disease and hypertensive nephropathy. Understanding the roles and regulatory mechanisms of these proteins in podocytes can provide valuable insights into the pathophysiology of kidney diseases and identify potential targets for therapeutic intervention.

Furthermore, our findings indicate that advanced mechanical stretch results in higher number of altered proteins. This observation is consistent with the established correlation between elevated blood pressure levels and pronounced glomerular alterations.^52–54^ Among the regulated proteins were numerous proteins with a function in RNA splicing, such as YB-1, which is translocated into the nucleus for transcriptional regulation under cellular stress conditions, including cold shock and oxidative stress.^55,56^ This demonstrates the potential significance of alternative splicing for the adaptation of podocytes to mechanical stress.

In podocytes, alternative splicing seems to affect several key genes involved in cytoskeletal dynamics, cell signaling, and adhesion. For instance, the alternative splicing of nephrin, a critical slit diaphragm protein, can influence its interaction with other podocyte proteins and the structural integrity of the filtration barrier.^57,58^ Similarly, splicing variations in the WT1 gene, a well-known regulator of podocyte differentiation, produce different isoforms that can modulate gene expression patterns crucial for podocyte differentiation and stress.^59–61^

In the present study, we have identified numerous novel genes that undergo alternative splicing in response to mechanical stretch. The most prevalent AS event identified in our study was exon skipping, a finding that is supported by data from Sugnet *et al.*.^62^

A comprehensive analysis of alternative splicing was conducted to assess the relative strengths and limitations of various computational systems, including rMATS, leafCutter, and Whippet. Some models demonstrated high precision and robustness in identifying splicing events, particularly in the presence of biological replicates. Conversely, others were designed to discover new events without the need for pre-defined reference annotations.^38–40^ Here we used several tools to increase the robustness of AS events detection. Consequently, we also used the IsoformSwitchAnalyzeR algorithm which is designed particularly to detect isoforms with opposite regulation .^37^

The robustness of our findings is evidenced by the observation that our most promising candidates (Myl6 and Shroom3) were identified as alternatively spliced in all utilized AS tools. Furthermore, both were expressed in podocytes and showed a significant isoform switch after mechanical stretch.

MYL6, being part of the actin-myosin complex, contributes to the organization and function of the actin cytoskeleton in podocytes, which is critical for maintaining the structural integrity of podocytes. In podocytes, the expression of *Myl6* is markedly high; this could indicate that *Myl6* plays an important role in the adaptation of cells to mechanical stretch and in the reorganization of cell structure under pathological conditions such as hypertension. By single-cell RNA sequencing and RNA FISH, the group of J. Salzman detected *Myl6* spliced isoforms in murine smooth muscle cells and human lung.^63^ To the best of our knowledge, we are the first to detect the existence of two distinct alternatively spliced Myl6 isoforms in podocytes. In glomerular diseases that damage podocytes, such as hypertensive nephropathy, dysregulation of *Myl6* splicing may impair the ability of podocytes to maintain their structure and function.

SHROOM3 is a crucial protein in podocytes and is involved in maintaining the cytoskeletal architecture of podocytes.^27^ Furthermore, genome-wide association studies have found that SNPs in *SHROOM3* are associated with chronic kidney disease (CKD) and lower estimated Glomerular Filtration Rate (eGFR).^22–26^ It interacts with actin filaments to regulate cell shape and adhesion, ensuring that podocytes can withstand the mechanical stress. Disruption in *Shroom3* expression or function can lead to cytoskeletal disorganization, resulting in podocyte effacement and proteinuria, a hallmark of glomerular diseases.^28^ Our study represents the inaugural investigation to demonstrate that specific *Shroom3* transcript isoforms are exclusively expressed in podocytes and that their expression levels undergo significant alterations in various injury models. This knowledge could be used to utilize specific *Shroom3* isoforms as splicing biomarkers for the diagnosis of patients with hypertensive nephropathy. The importance of *Shroom3* for podocytes was impressively demonstrated by knockdown experiments, which showed the essential role of *Shroom3* for the stability and maintenance of the actin cytoskeleton in podocytes. The loss of *Shroom3* appears to result in an impaired polymerization of G-actin monomers to F-actin. This finding is consistent with previous reports indicating that all Shroom family proteins interact with actin and are essential for actin polymerization.^64^

It is noteworthy that there is a link between SHROOM3 and MYL6 (as part of the essential light chain of myosin II).^65–67^ It is known that the ASD2 domain of Shroom3 can bind directly to Rho-associated kinases (ROCKs).^68–70^ Once ROCKs are activated by RhoA, they phosphorylate the myosin regulatory light chain to activate myosin II, resulting in F-actin contraction.^69,71–74^ This molecular pathway is supported by studies demonstrating that Shroom3 colocalizes with actin filaments, RhoA, ROCKs, and myosin II at adherens junctions.^69,75,76^ In Shroom3 homozygous null mutant mice podocytes, a reduced ROCK1 expression and phosphorylation was observed, which further confirmed that Shroom3 regulates the actomyosin cytoskeleton through the ROCK/Myosin II signaling pathways.^27^

In summary, understanding the role of alternative splicing in podocyte biology offers potential therapeutic avenues for hypertensive nephropathy. Targeting pathways involved in the regulation of critical podocyte genes, such as *Shroom3* and *Myl6*, could help restore normal splicing patterns and improve podocyte function. Small molecules or antisense oligonucleotides that modulate splicing could correct mis-splicing events and enhance podocyte resilience to hypertensive stress. Moreover, *Myl6* or *Shroom3* may serve as splicing biomarkers associated with hypertensive nephropathy, offering the potential for early diagnosis and personalized treatment strategies.

## Acknowledgements

The authors thank Anja Wiechert, Claudia Weber, and Sophia-Marie Bach for technical assistance.

This work was supported by grants of the Federal Ministry of Education and Research (BMBF, grant 01ZX1908B + 01ZX2208B, Sys_CARE) to Jan Baumbach, Uwe Völker, and Nicole Endlich. This work was generously supported by the Südmeyer fund for kidney and vascular research (‘Südmeyer Stiftung für Nieren-und Gefäßforschung’) and the Dr. Gerhard Büchtemann fund, Hamburg, Germany. The funders had no role in study design, data collection and analysis, decision to publish, or preparation of the manuscript.

## Author Contributions Statement

The study was designed by K.E., N.E. and F.K.; F.K., F.M. contributed to the cell culture experiments; biopsies were handled and analyzed by K.A., F.K., F.M.; LC-MS/MS was performed by E.H. and U.V.; RNA-Seq was performed, supervised and analyzed by O.T., C.L., T.K. and M.L.; protein structure analysis were done by A.G. and O.K., mice experiments were performed by L.D., U.W., F.M.; all other experiments were performed by F.K. and F.M.; experimental data were analyzed by F.K., F.M., O.T., S.A., S.S., C.L.; F.K., F.M. and N.E. wrote the main manuscript text. F.K. and F.M. prepared figures. All authors reviewed the manuscript.

## Data availability

All data supporting the findings of this study are available within the paper and its Supplementary Information.

The mass spectrometry proteomics data are available in supplementary data files (Supplementary Data 1) and have been deposited to the ProteomeXchange Consortium via the PRIDE partner repository with the dataset identifier PXD055066 and PXD055232 (reviewer login details for access available on request).

RNA-seq data are available in Supplementary Data 2. All other relevant data and materials are available from the corresponding author on reasonable request.

The RNA-Seq used in the study data (the raw data and the preprocessed data) has been shared privately via the GEO database. The reviewers can access the data using the reviewer token: https://www.ncbi.nlm.nih.gov/geo/query/acc.cgi?acc=GSE276267 (secure token for access by reviewer available on request).

The pipeline for analyzing RNA-Seq data is published via GitHub and can be reviewed: https://github.com/OlgaVT/Alternative-splicing-in-mechanically-stretched-podocytes-as-a-model-of-glomerular-hypertension.

---

**Fig. S1:**
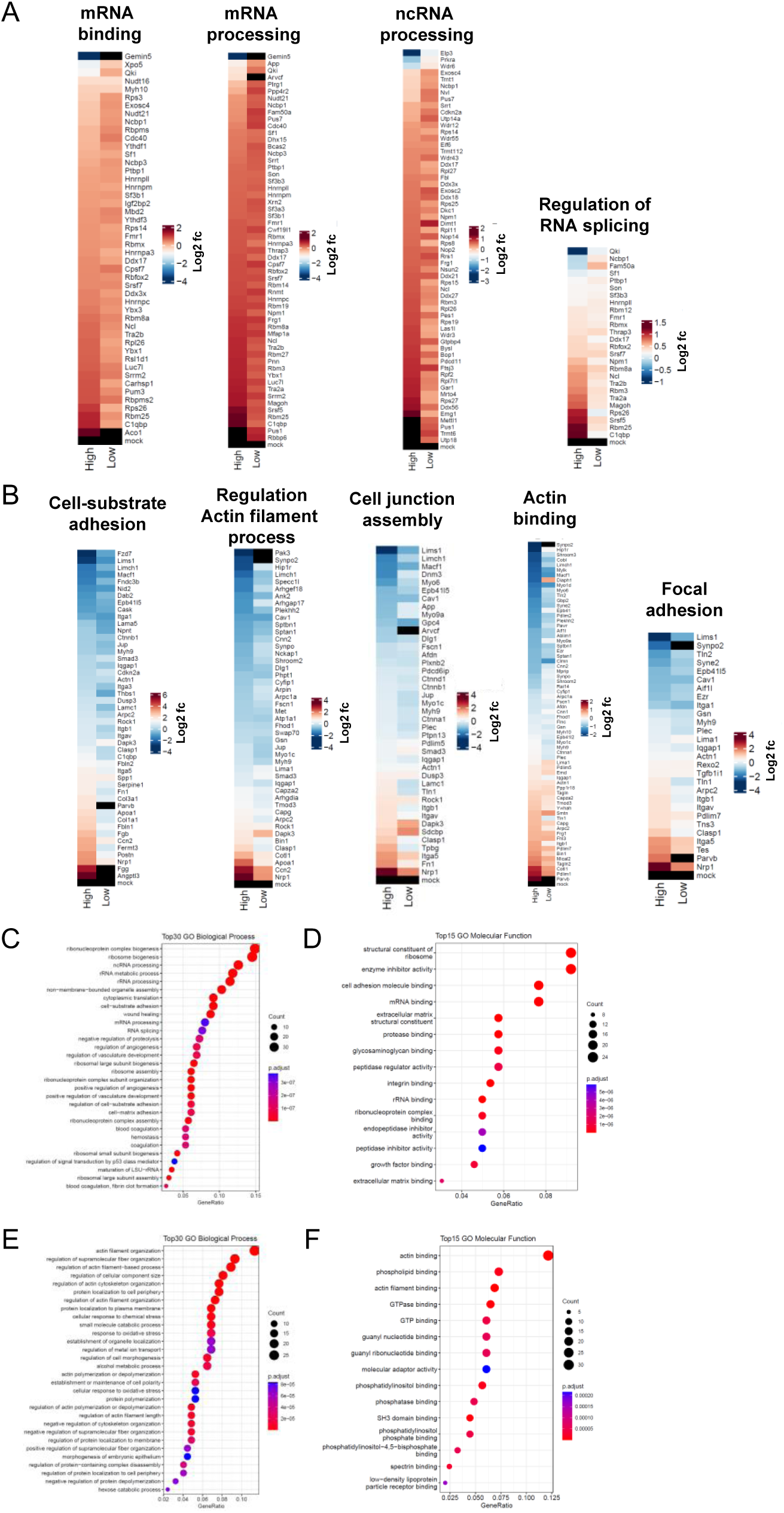
GO and GSEA from mechanically stretched podocytes (proteome based). Heatmap of GO terms of (A) up-regulated and (B) down-regulated proteins in high stretched podocytes. Data are represented as log2 fold change, a p-value < 0.05 was considered as significant. Red: up-regulated; blue: down-regulated compared to controls. (C-F) GO (Gene Ontology) clusters (Biological Process and Molecular Function) of low stretched podocytes.

**Fig. S2:**
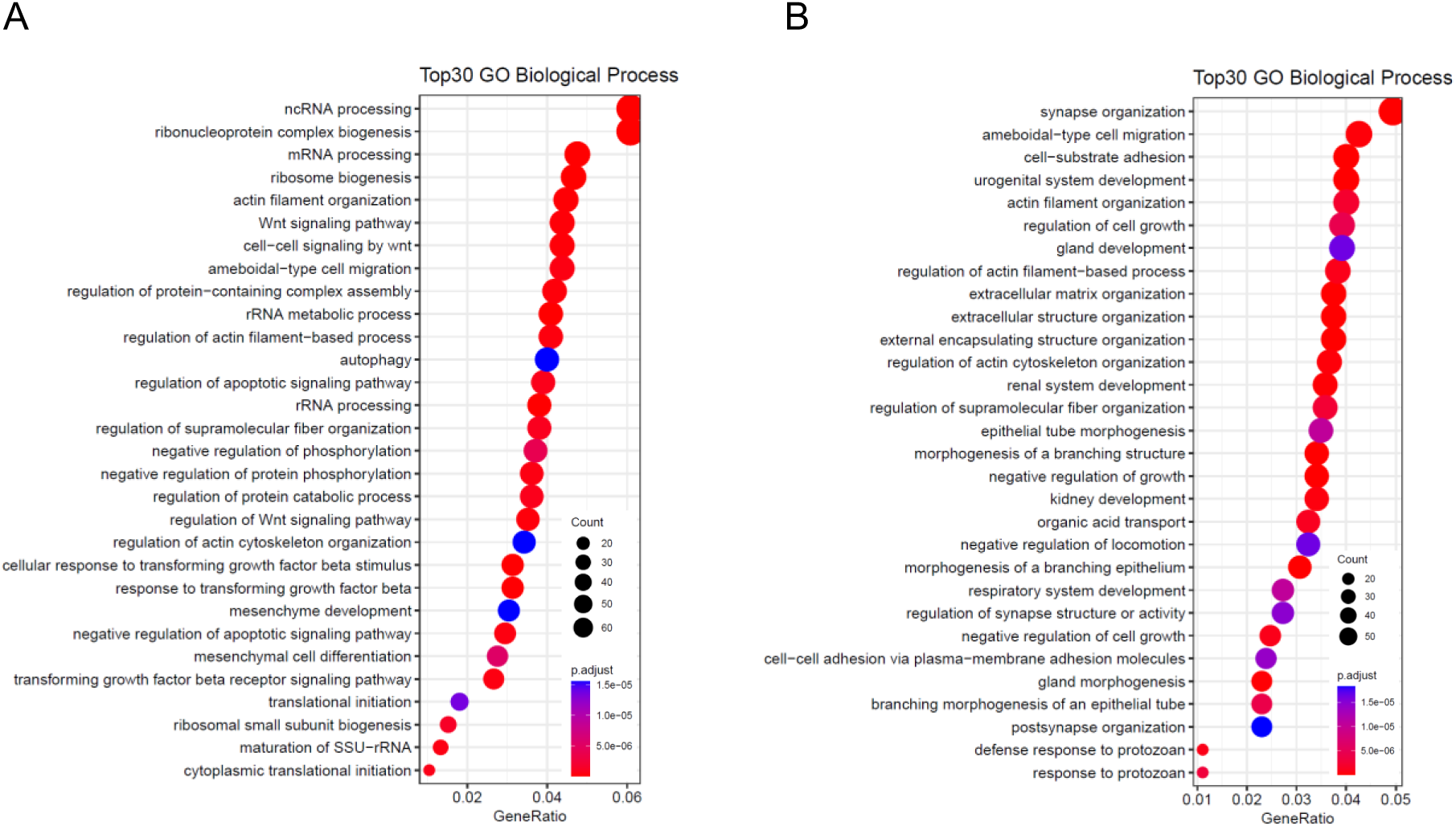
Detailed gene set enrichment analysis (GSEA) of significantly regulated genes (transcriptome based). RNA-Seq based enriched Top30 GO (Gene Ontology) clusters (Biological Process) of mechanically stretched podocytes. Demonstrated are GO clusters from differentially up-regulated transcripts (A) and down-regulated transcripts (B).

**Fig. S3:**
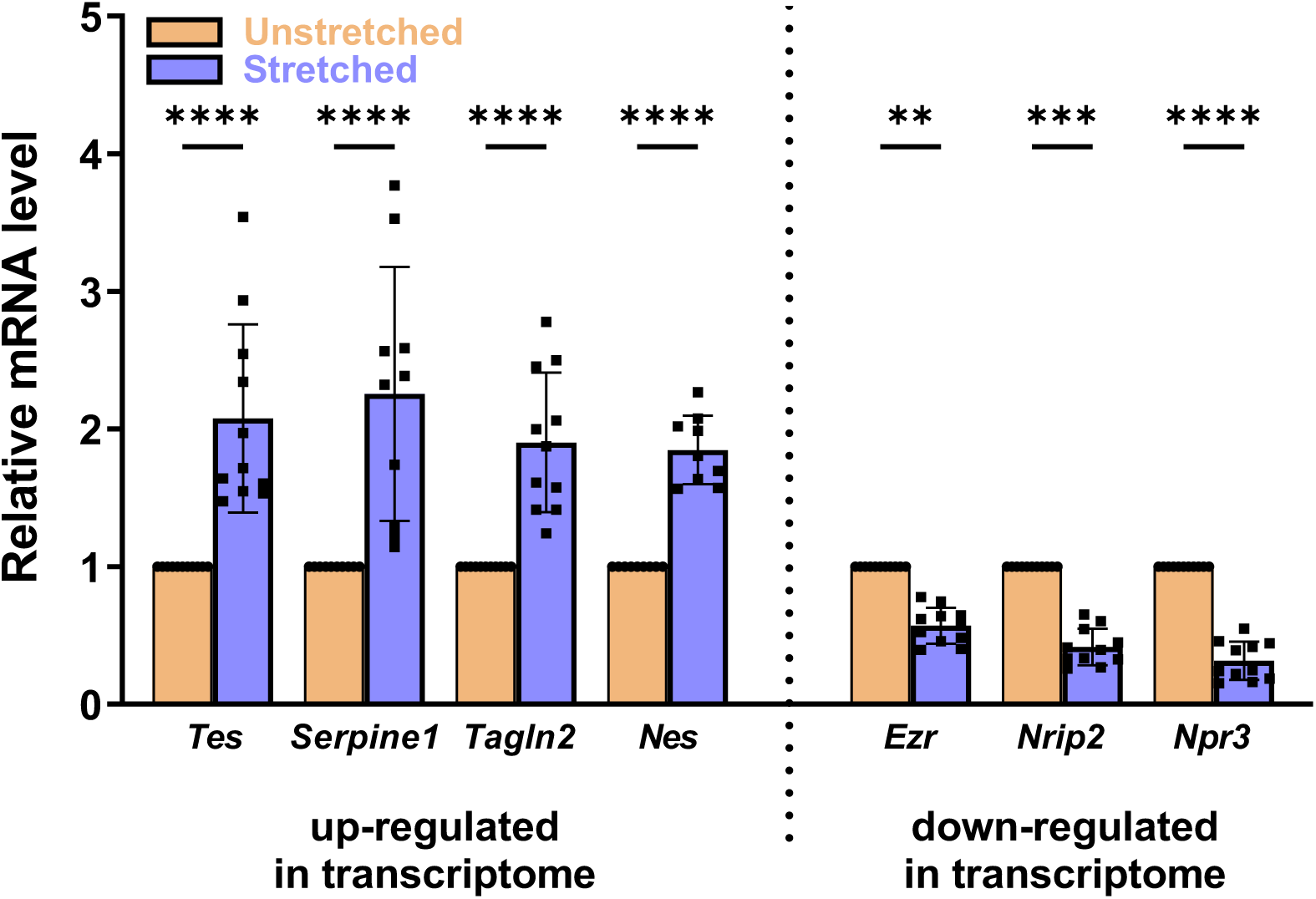
Verification of up- and down-regulated candidates from Fig. 2. (A) The mRNA expression of *Tes*, *Serpine11*, *Tagln2*, *Nes*, *Ezr*, *Nrip2,* and *Npr3* in stretched and unstretched podocytes was quantified by qRT-PCR (n≥9). qRT-PCR experiments were normalized to the unstretched samples, *Gapdh* served as a reference. Data are presented as means ± SD. ** *p*<0.01; *** *p*<0.001; **** *p*<0.0001.

**Fig. S4:**
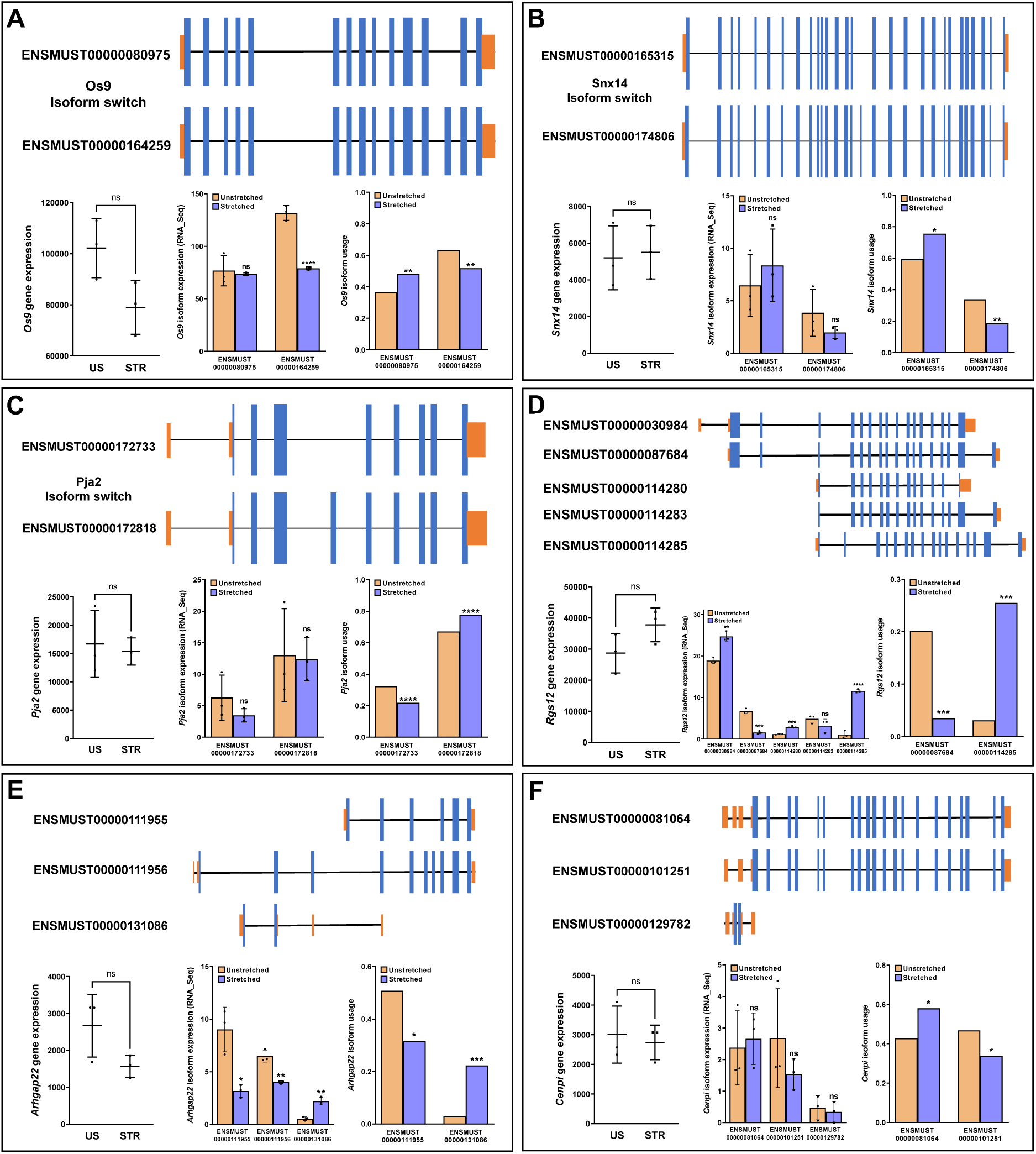
Isoform switch detection of Os9, Snx14, Pja2, Rgs12, Arhgap22 and Cenpi. Schematic overview of A) *Os9*, B) *Snx14*, C) *Pja2*, D) *Rgs12*, E) *Arhgap22* and F) *Cenpi* isoforms found in murine cultured podocytes (Blue: coding exons; Orange: untranslated regions). Each partial figure displays the corresponding gene and transcript expression in unstretched (US) and mechanically stretched (STR) podocytes, as well as isoform usage based on IsoformSwitchAnalyzer data. Data are presented as means ± SD. * *p*<0.05; ** *p*<0.01; *** *p*<0.001; **** *p*<0.0001; ns, not significant.

**Fig. S5:**
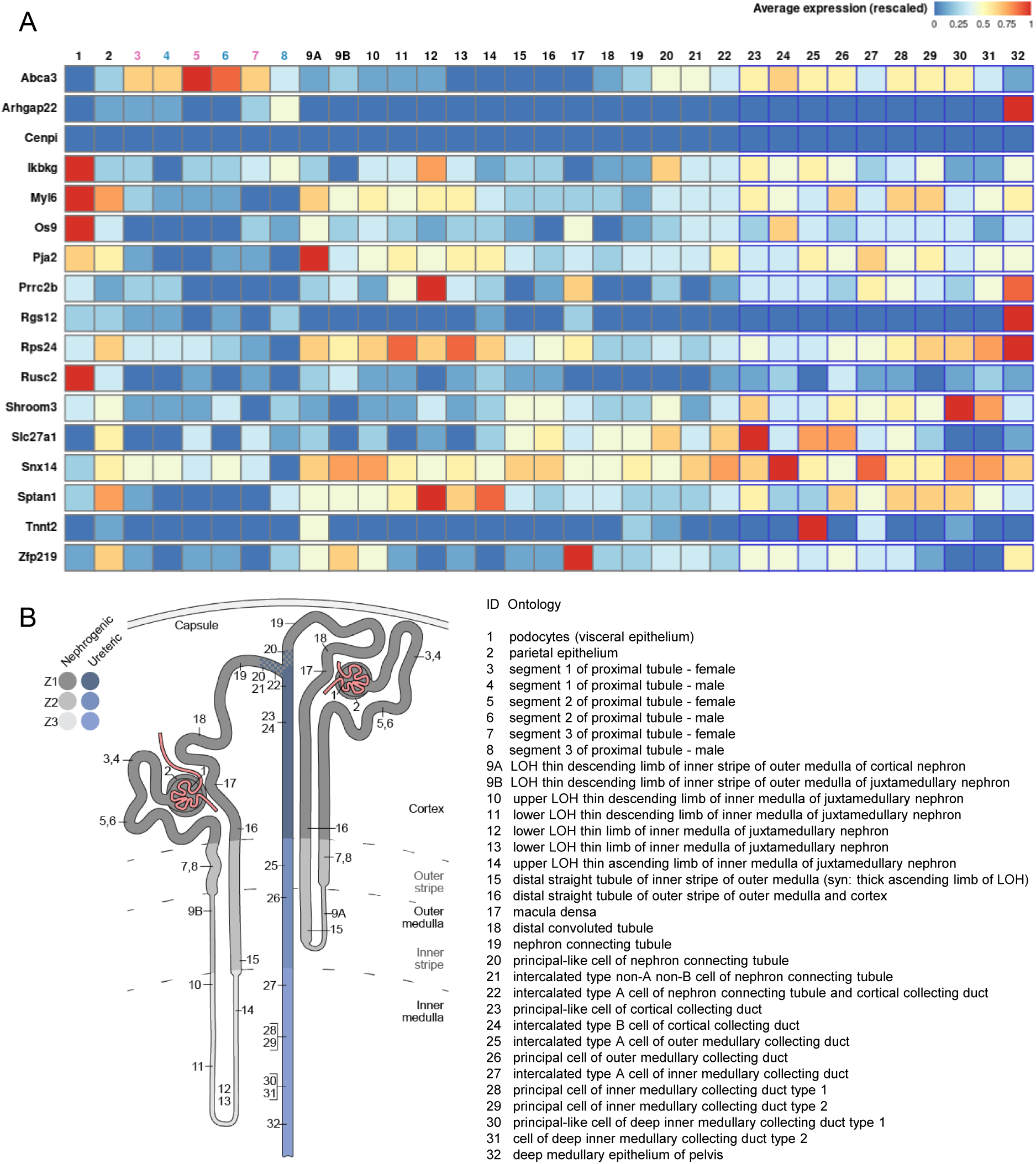
Kidney expression pattern of RNA-Seq based alternative spliced candidates. (A) Database analysis using the Kidney Cell Explorer by Ransick et al.^77^ based on a single-cell RNA sequencing data set of murine kidneys showed the expression pattern for all kidney cell fractions. Color code: Red means a high average expression; Blue: low expression. (B) Legend of the numerical labeling of figure A.

**Fig. S6:**
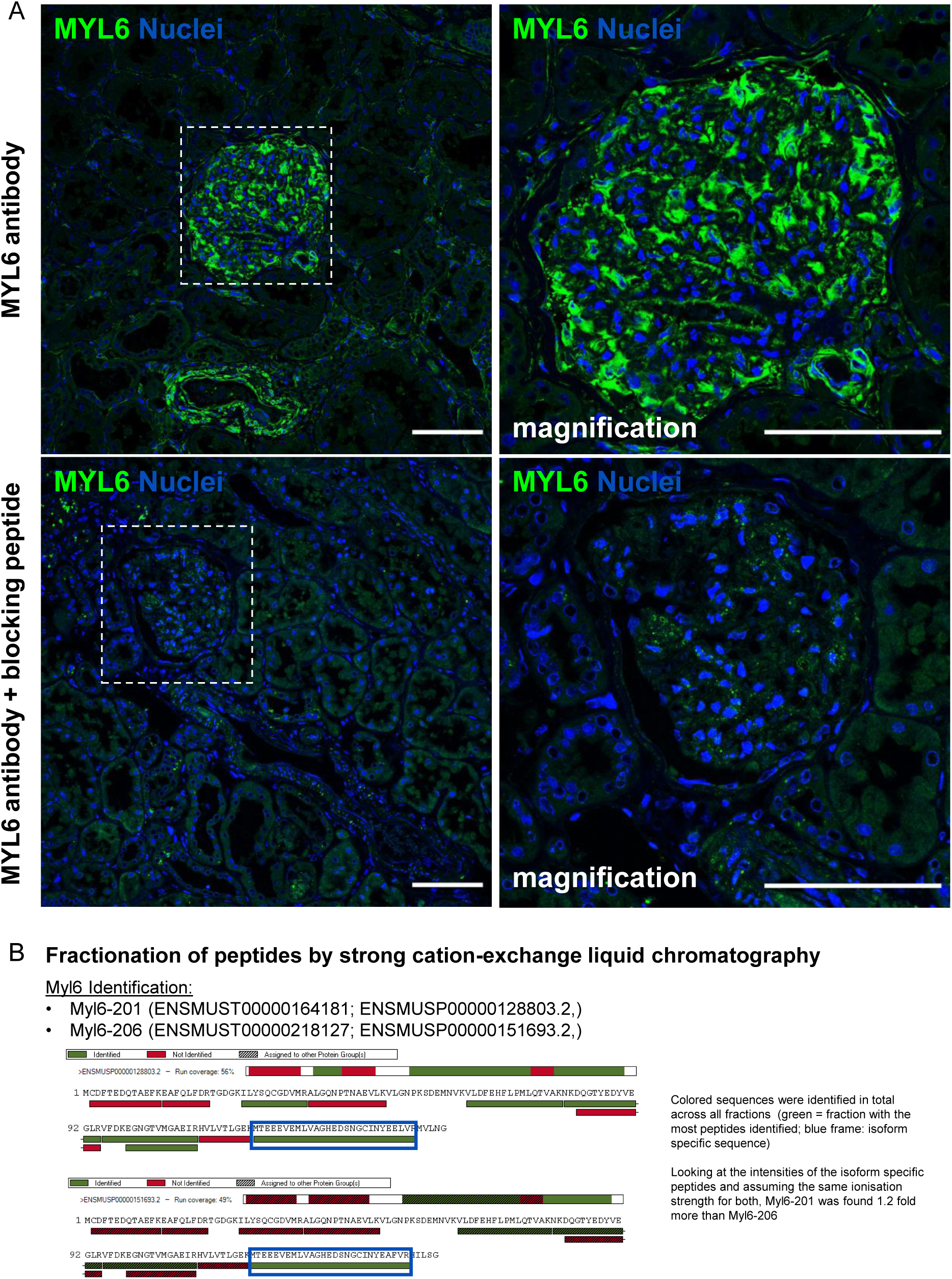
MYL6 antibody verification and confirmation of two Myl6 isoforms in podocytes by strong cation exchange (SCX) chromatography. (A) Immunofluorescence staining with anti-MYL6 antibody (shown in green) in mouse kidney tissue. The specificity of the MYL6 antibody was successfully demonstrated by the use of a MYL6 blocking peptide (lower panel). Nuclei were stained with HOECHST (blue). Scale bars represent 50 µm. (B) Fractionation of Myl6 peptides by strong cation-exchange liquid chromatography confirmed the presence of the two Myl6 isoforms Myl6-201 and Myl6-206.

**Fig. S7:**
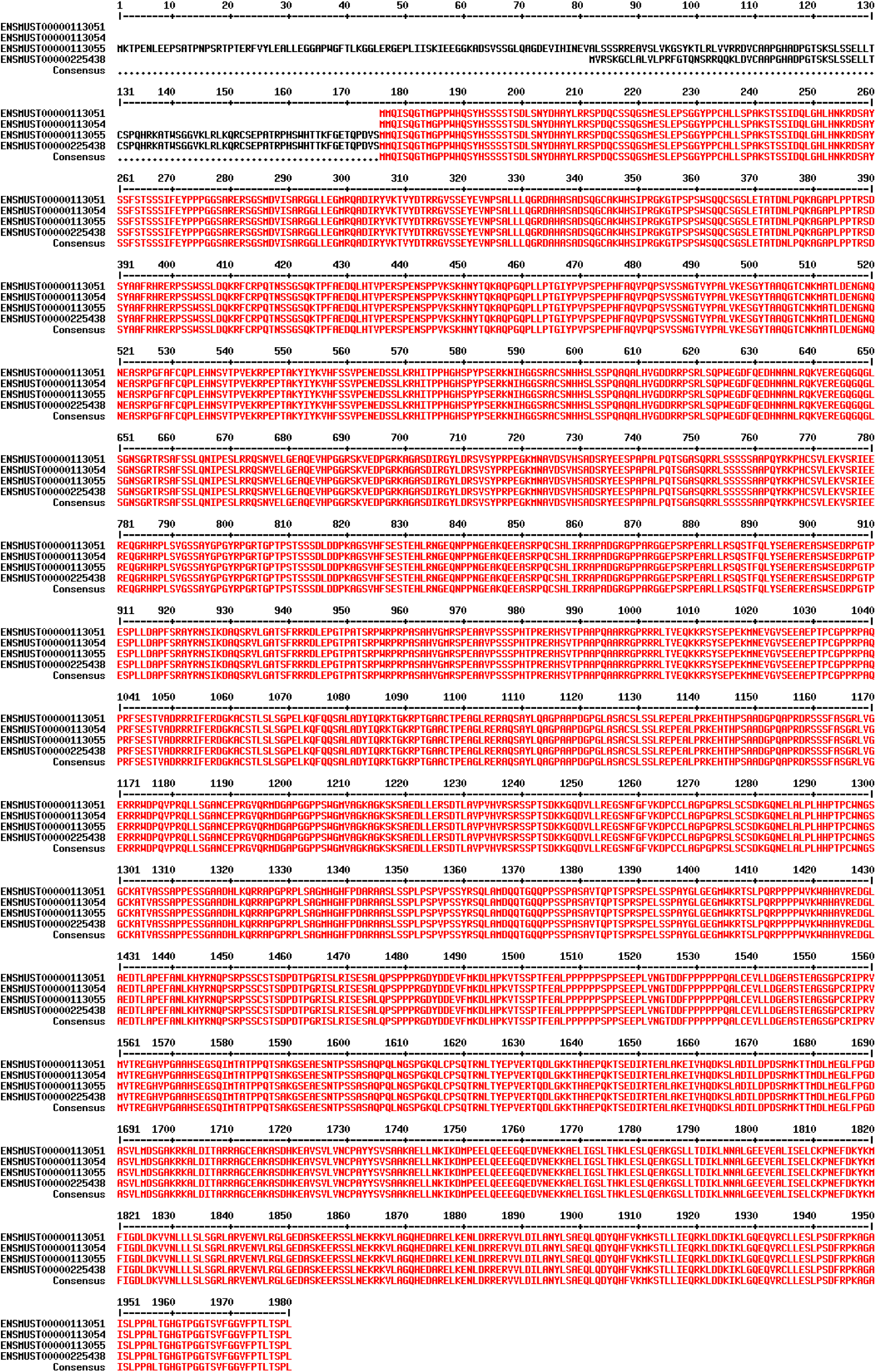
Mouse SHROOM3 protein sequence. Protein sequence alignment of all murine SHROOM3 isoforms.

**Fig. S8:**
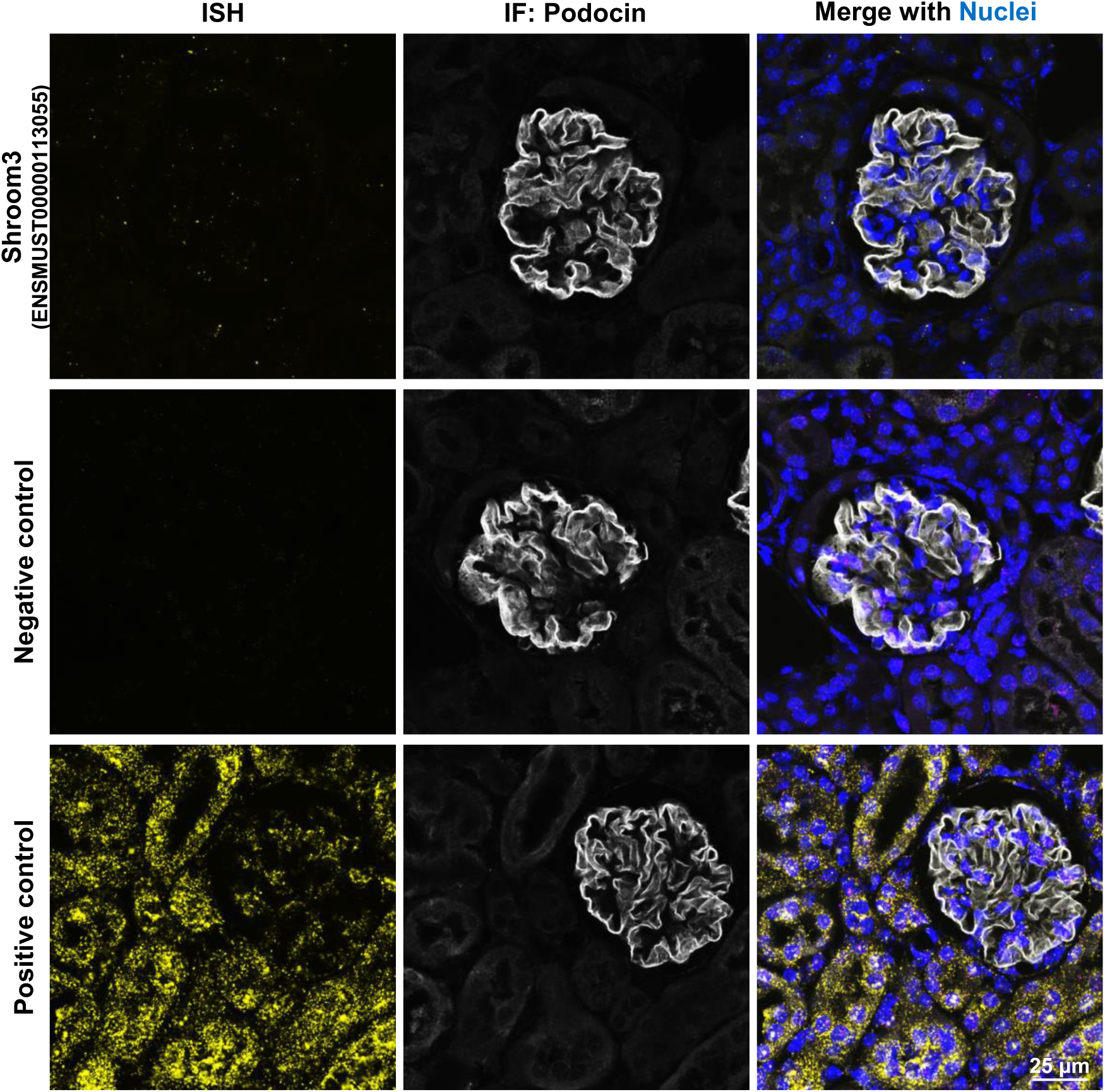
RNAscope^®^ ISH Assay. *In situ* hybridization of the *Shroom3* coded isoform ENSMUST00000113055 in mouse kidney (yellow dots). Immunostaining of podocin is shown in white. Nuclei stained with HOECHST (blue). The negative control (*dapB*) and the positive control (*Ppib*) are also shown. Scale bar represents 25 μm.

**Fig. S9:**
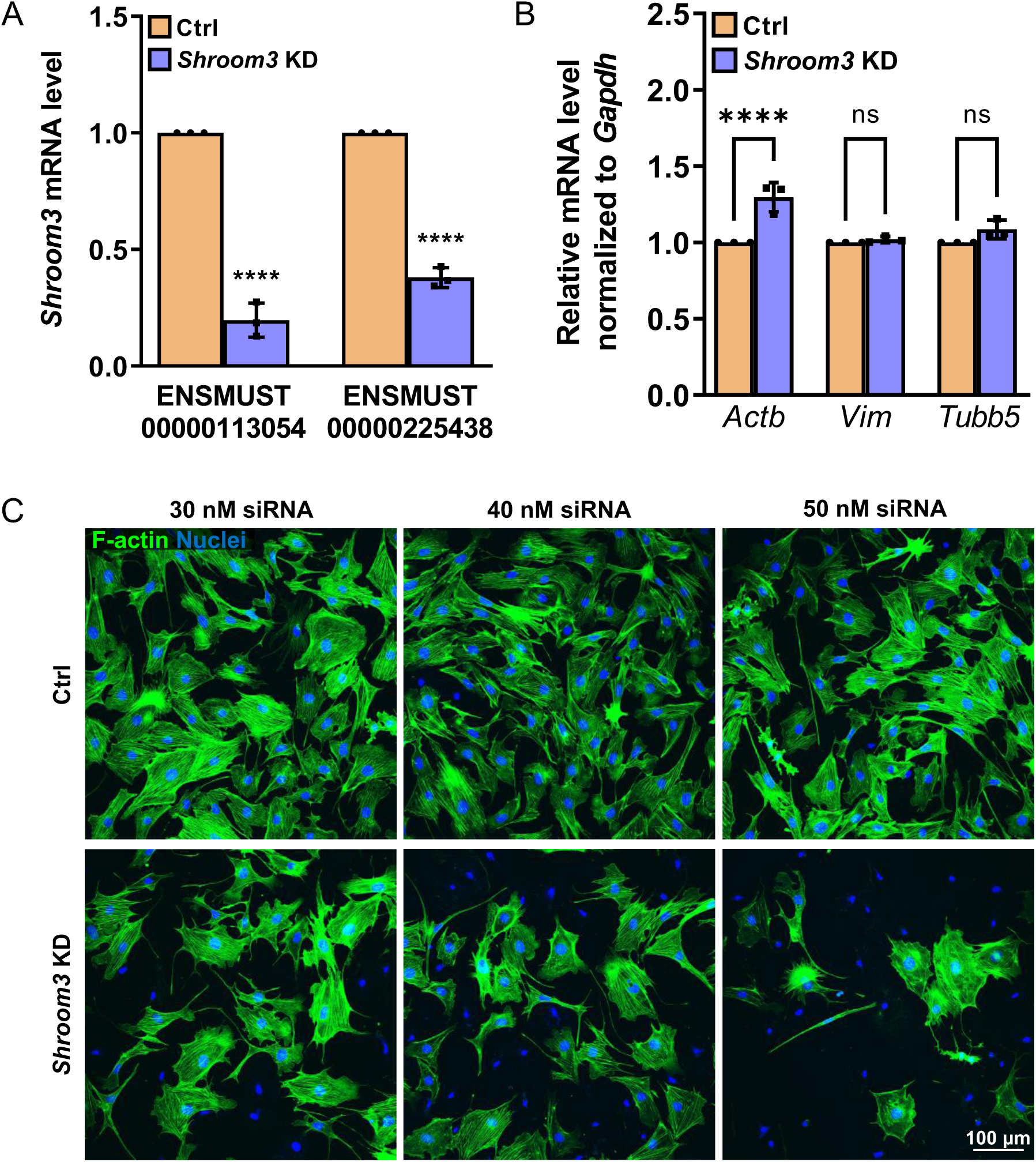
*Shroom3* Knockdown (KD) in cultured murine podocytes. (A) qRT-PCR quantification of *Shroom3* knockdown (*Shroom3* KD) podocytes showed a significant decrease of *Shroom3* coded transcript expression levels compared to the control (Ctrl). (B) Beta-Actin (*Actb*) mRNA level was significantly up-regulated in Shroom3 KD podocytes. In contrast, the mRNA level of vimentin (*Vim*) as a marker of intermediate filaments and β-tubulin (*Tubb5*) were not changed significantly. Data are normalized against Ctrl KD and Gapdh and presented as means ± SD. (C) With an increased siRNA concentration the number of cells without visible F-actin increased dramatically in *Shroom3* KD podocytes. F-Actin is shown in green. Nuclei were stained with DAPI (blue). Scale bar represents 100 μm. ** *p*<0.01; **** *p*<0.0001.

