## Supplementary information for "Alternative splicing in mechanically stretched podocytes as a model of glomerular hypertension"

#### Supplementary Figures

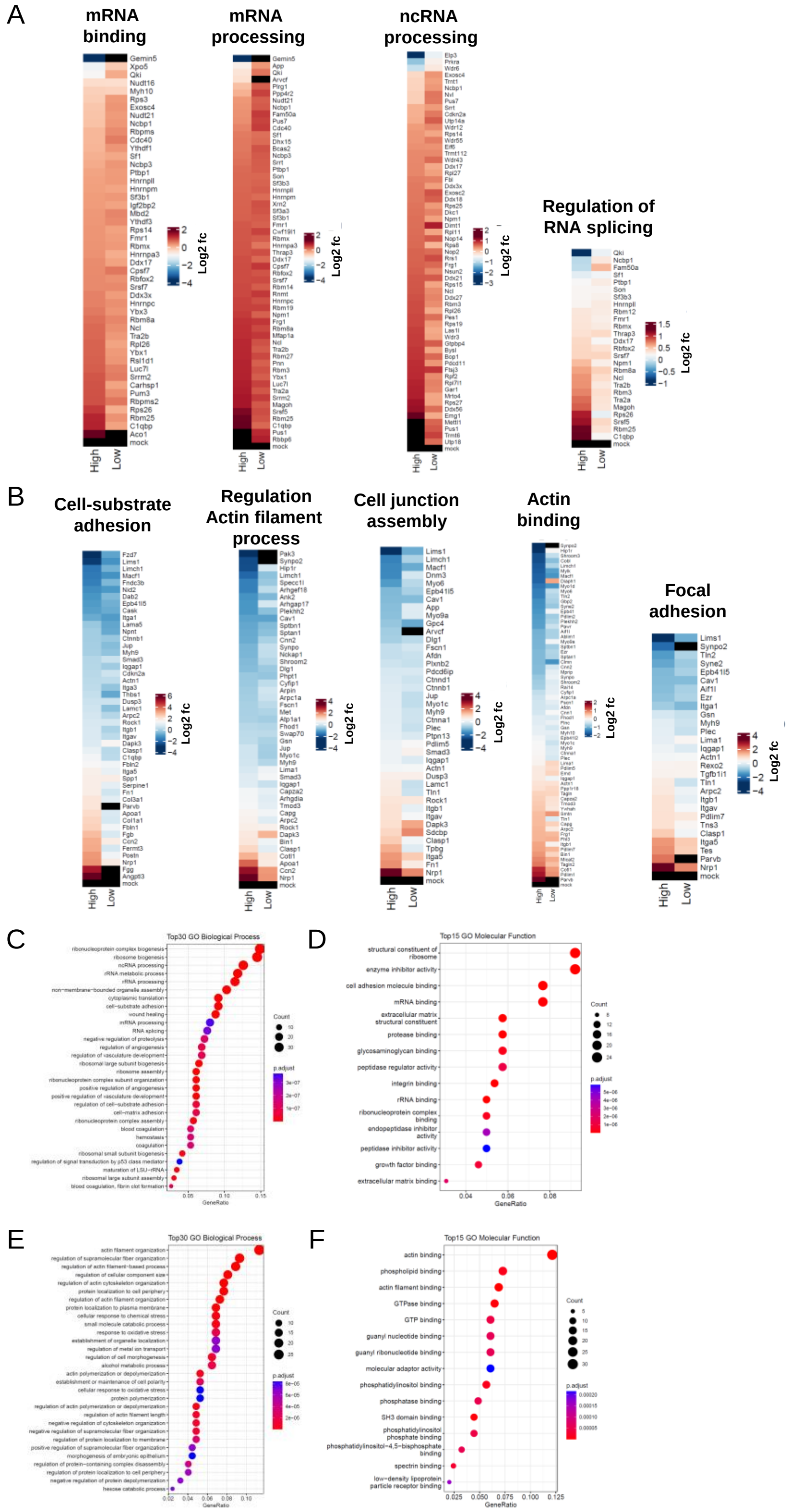

**Fig. S1: GO and GSEA from mechanically stretched podocytes (proteome based).**
Heatmap of GO terms of (A) up-regulated and (B) down-regulated proteins in high stretched podocytes. Data are represented as log2 fold change, a p-value < 0.05 was considered as significant. Red: up-regulated; blue: down-regulated compared to controls. (C-F) GO (Gene Ontology) clusters (Biological Process and Molecular Function) of low stretched podocytes.

A B

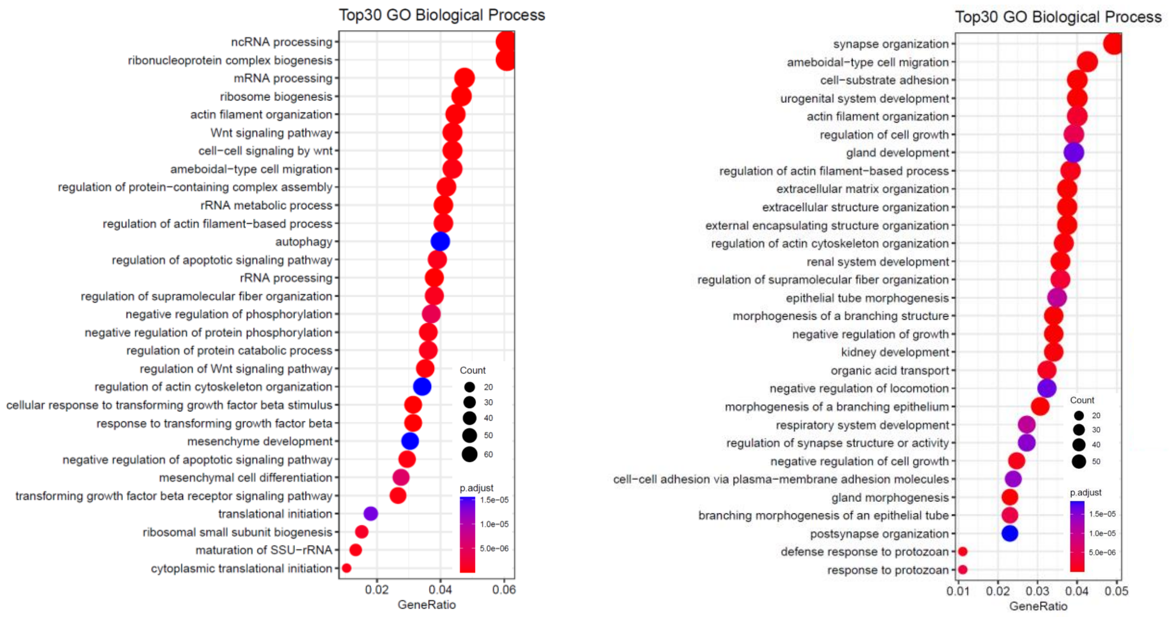

**Fig. S2: Detailed gene set enrichment analysis (GSEA) of significantly regulated genes (transcriptome based).**

RNA-Seq based enriched Top30 GO (Gene Ontology) clusters (Biological Process) of mechanically stretched podocytes. Demonstrated are GO clusters from differentially up-regulated transcripts (A) and down-regulated transcripts (B).

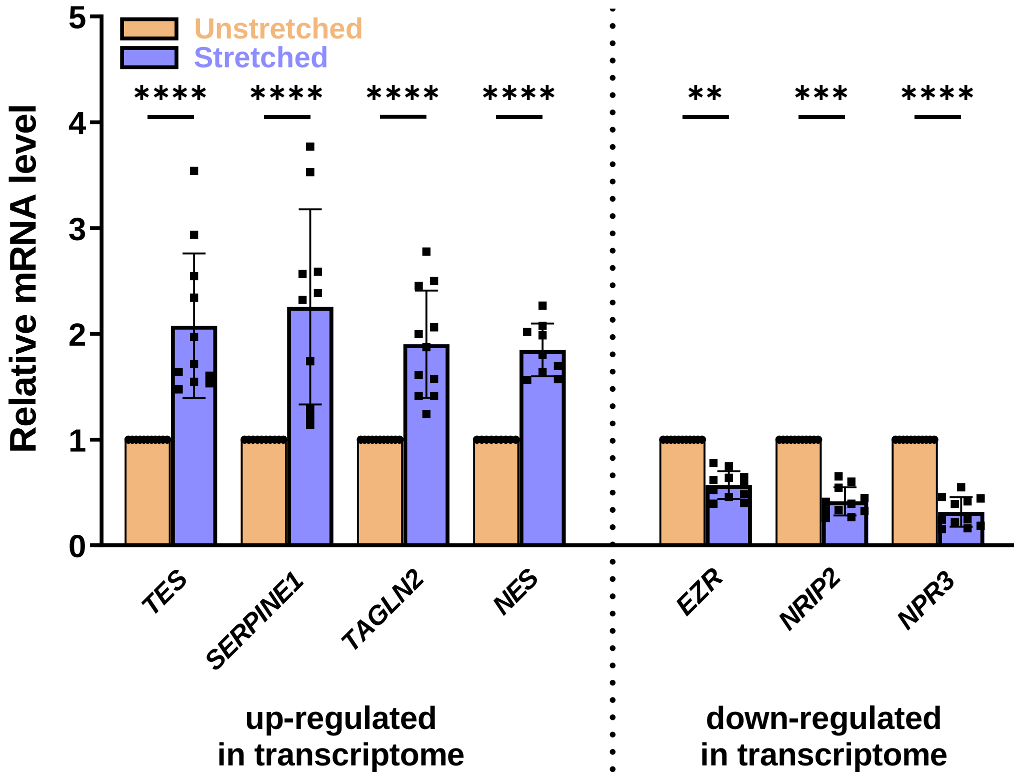

**Fig. S3: Verification of up- and down-regulated candidates from Fig. 2.**
(A) The mRNA expression of *Tes*, *Serpine11*, *Tagln2*, *Nes*, *Ezr*, *Nrip2,* and *Npr3* in stretched and unstretched podocytes was quantified by qRT-PCR (n≥9). qRT-PCR experiments were normalized to the unstretched samples, *Gapdh* served as a reference. Data are presented as means ± SD. ** $p<0.01$; *** $p<0.001$; **** $p<0.0001$.

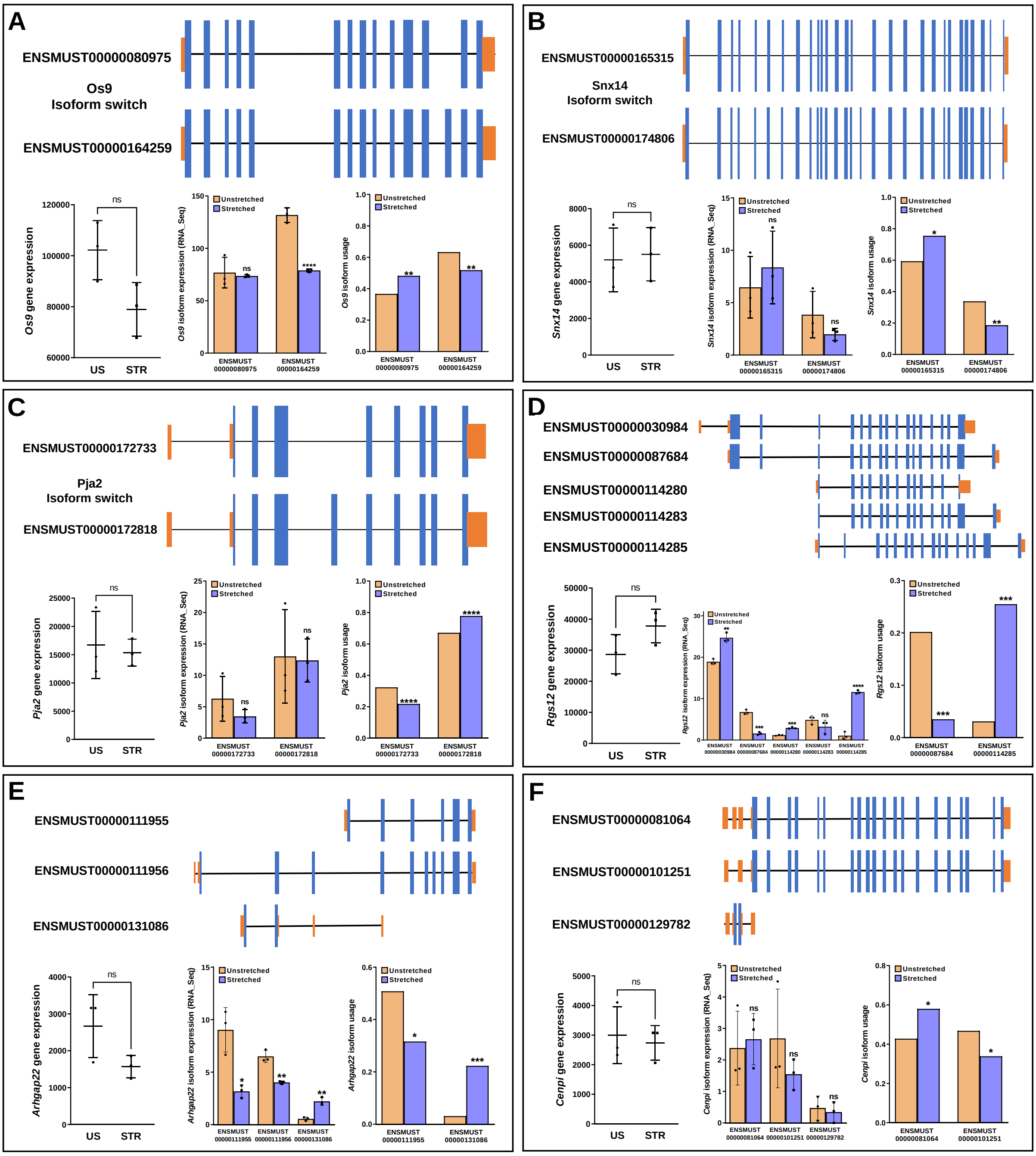

**Fig. S4: Isoform switch detection of Os9, Snx14, Pja2, Rgs12, Arhgap22 and Cenpi.** Schematic overview of A) *Os9*, B) *Snx14*, C) *Pja2*, D) *Rgs12*, E) *Arhgap22* and F) *Cenpi* isoforms found in murine cultured podocytes (Blue: coding exons; Orange: untranslated regions). Each partial figure displays the corresponding gene and transcript expression in unstretched (US) and mechanically stretched (STR) podocytes, as well as isoform usage based on IsoformSwitchAnalyzer data. Data are presented as means ± SD. * $p$<0.05; ** $p$<0.01; *** $p$<0.001; **** $p$<0.0001; ns, not significant.

A

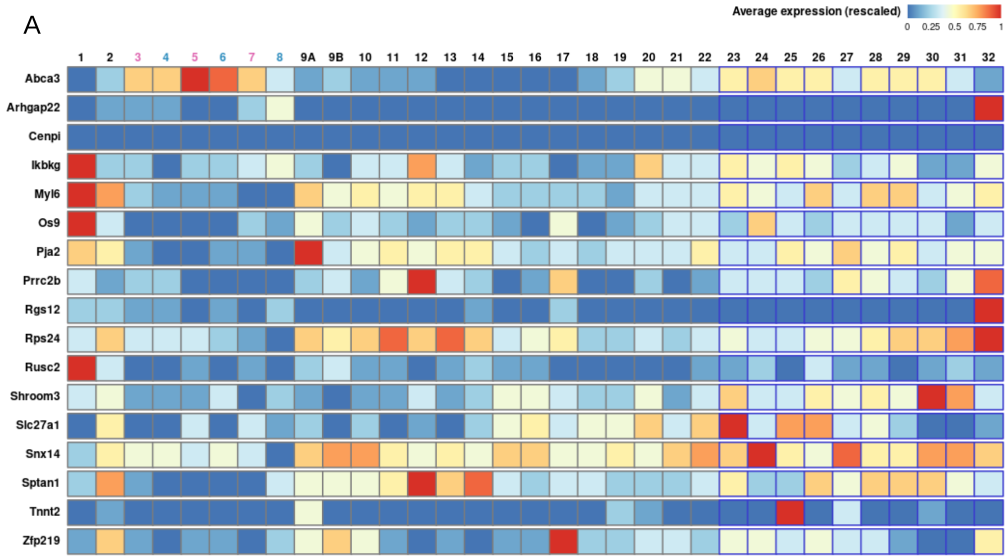

B

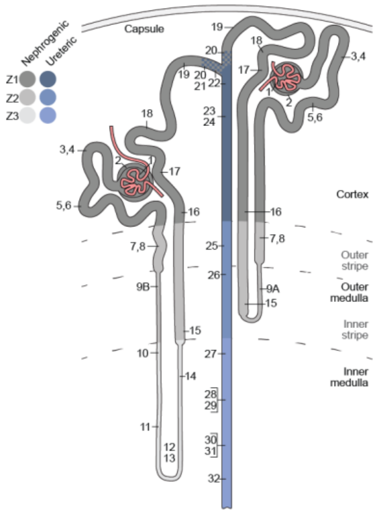

ID Ontology

1 podocytes (visceral epithelium)
2 parietal epithelium
3 segment 1 of proximal tubule - female
4 segment 1 of proximal tubule - male
5 segment 2 of proximal tubule - female
6 segment 2 of proximal tubule - male
7 segment 3 of proximal tubule - female
8 segment 3 of proximal tubule - male
9A LOH thin descending limb of inner stripe of outer medulla of cortical nephron
9B LOH thin descending limb of inner stripe of outer medulla of juxtamedullary nephron
10 upper LOH thin descending limb of inner medulla of juxtamedullary nephron
11 lower LOH thin descending limb of inner medulla of juxtamedullary nephron
12 lower LOH thin limb of inner medulla of juxtamedullary nephron
13 lower LOH thin limb of inner medulla of juxtamedullary nephron
14 upper LOH thin ascending limb of inner medulla of juxtamedullary nephron
15 distal straight tubule of inner stripe of outer medulla (syn: thick ascending limb of LOH)
16 distal straight tubule of outer stripe of outer medulla and cortex
17 macula densa
18 distal convoluted tubule
19 nephron connecting tubule
20 principal-like cell of nephron connecting tubule
21 intercalated type non-A non-B cell of nephron connecting tubule
22 intercalated type A cell of nephron connecting tubule and cortical collecting duct
23 principal-like cell of cortical collecting duct
24 intercalated type B cell of cortical collecting duct
25 intercalated type A cell of outer medullary collecting duct
26 principal cell of outer medullary collecting duct
27 intercalated type A cell of inner medullary collecting duct
28 principal cell of inner medullary collecting duct type 1
29 principal cell of inner medullary collecting duct type 2
30 principal-like cell of deep inner medullary collecting duct type 1
31 cell of deep inner medullary collecting duct type 2
32 deep medullary epithelium of pelvis

**Fig. S5: Kidney expression pattern of RNA-Seq based alternative spliced candidates.**
(A) Database analysis using the Kidney Cell Explorer by Ransick et al [PMID: 31689386] based on a single-cell RNA sequencing data set of murine kidneys showed the expression pattern for all kidney cell fractions. Color code: Red means a high average expression; Blue: low expression. (B) Legend of the numerical labeling of figure A.

A

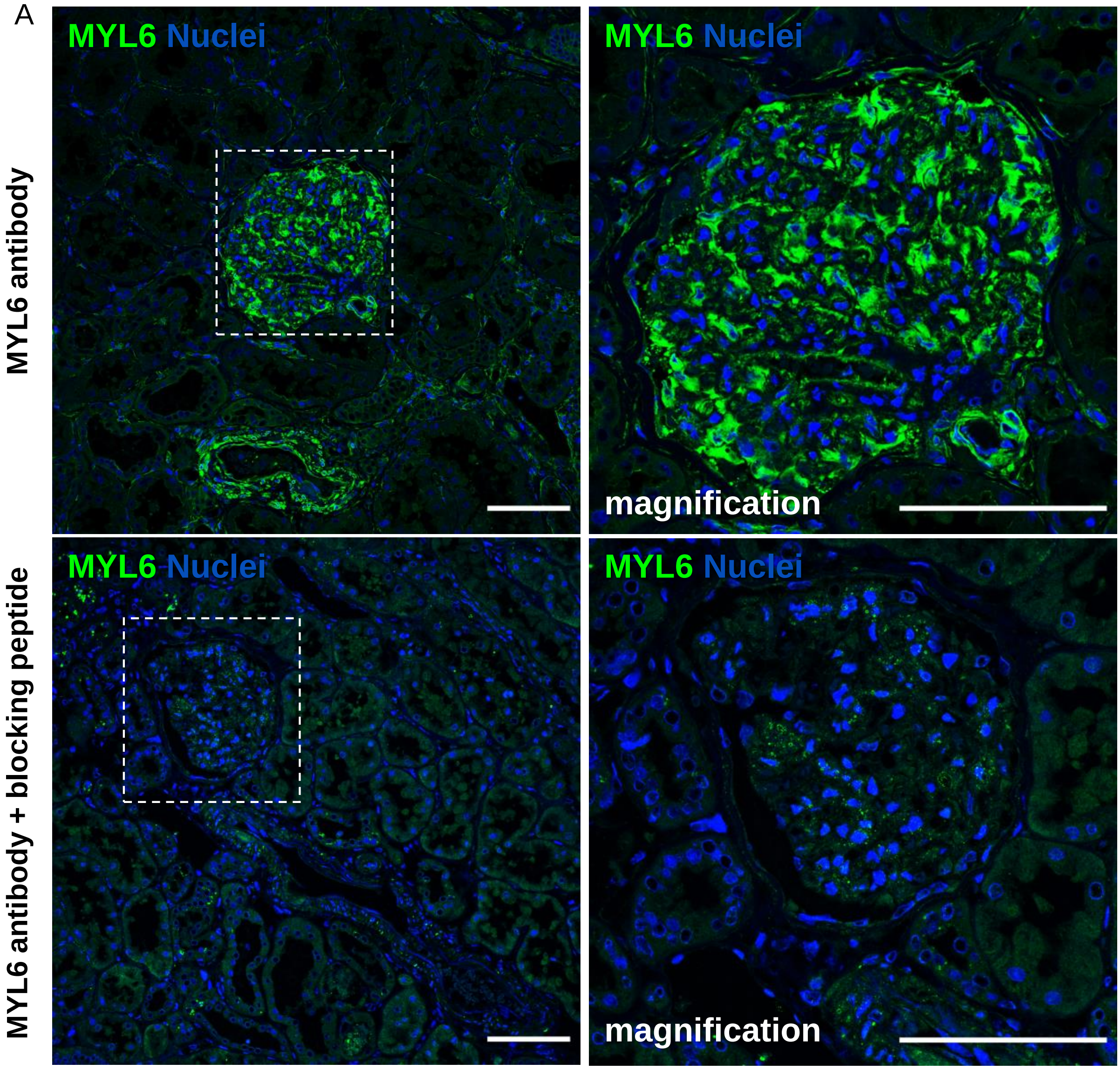

B **Fractionation of peptides by strong cation-exchange liquid chromatography**

Myl6 Identification:

- Myl6-201 (ENSMUST00000164181; ENSMUSP00000128803.2,)
- Myl6-206 (ENSMUST00000218127; ENSMUSP00000151693.2,)

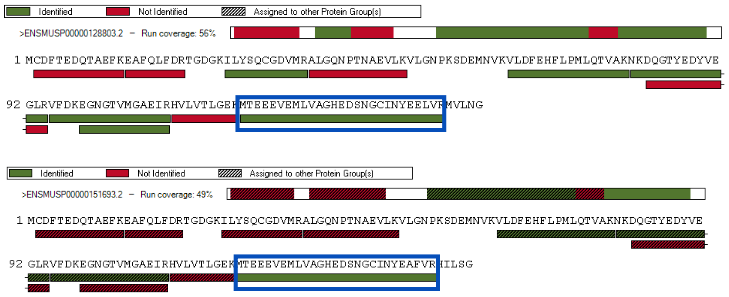

Colored sequences were identified in total across all fractions (green = fraction with the most peptides identified; blue frame: isoform specific sequence)

Looking at the intensities of the isoform specific peptides and assuming the same ionisation strength for both, Myl6-201 was found 1.2 fold more than Myl6-206

**Fig. S6: MYL6 antibody verification and confirmation of two Myl6 isoforms in podocytes by strong cation exchange (SCX) chromatography.**
(A) Immunofluorescence staining with anti-MYL6 antibody (shown in green) in mouse kidney tissue. The specificity of the MYL6 antibody was successfully demonstrated by the use of a MYL6 blocking peptide (lower panel). Nuclei were stained with HOECHST (blue). Scale bars represent 50 µm. (B) Fractionation of Myl6 peptides by strong cation-exchange liquid chromatography confirmed the presence of the two Myl6 isoforms Myl6-201 and Myl6-206.

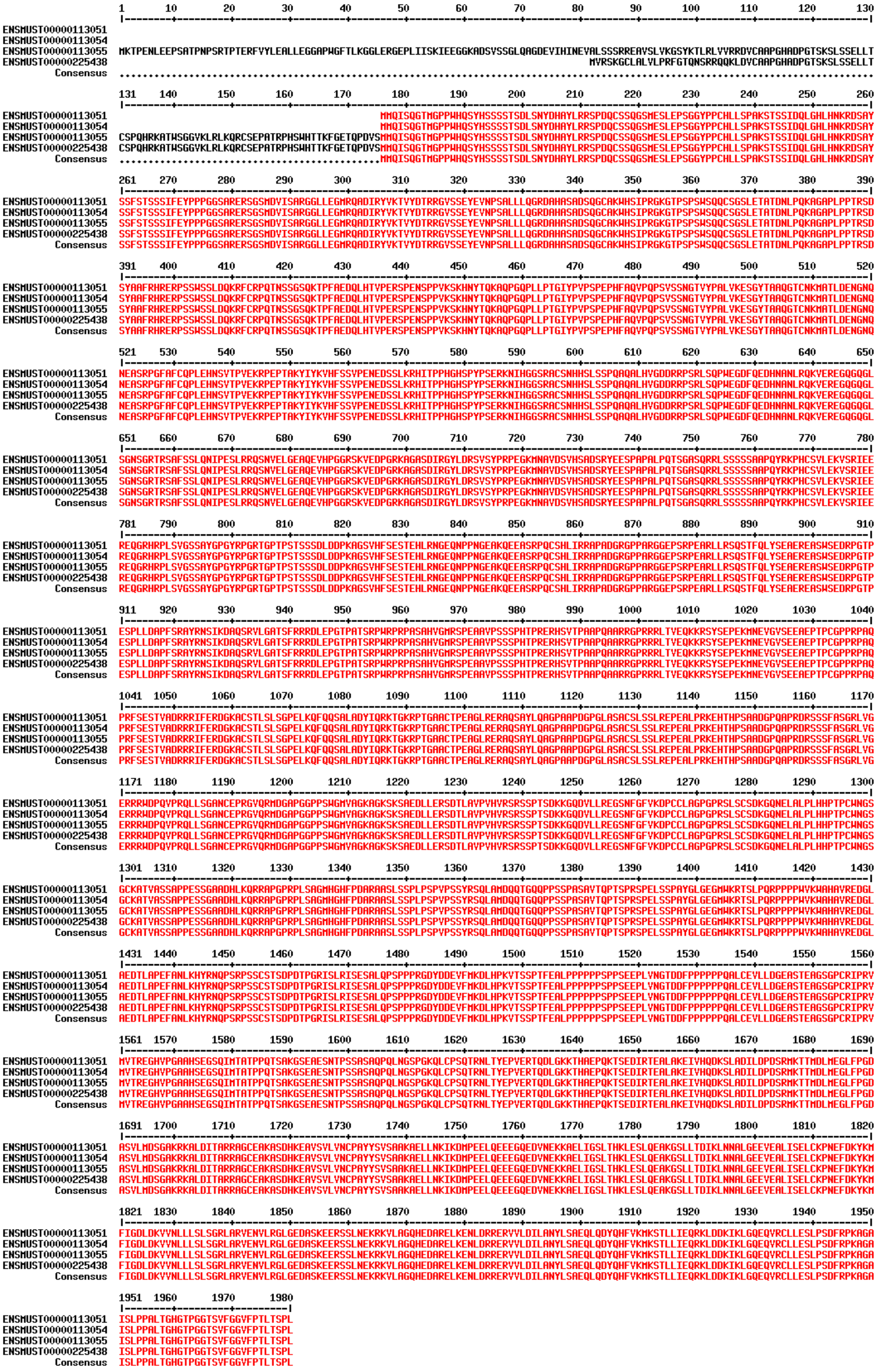

**Fig. S7: Mouse SHROOM3 protein sequence.**

Protein sequence alignment of all murine SHROOM3 isoforms.

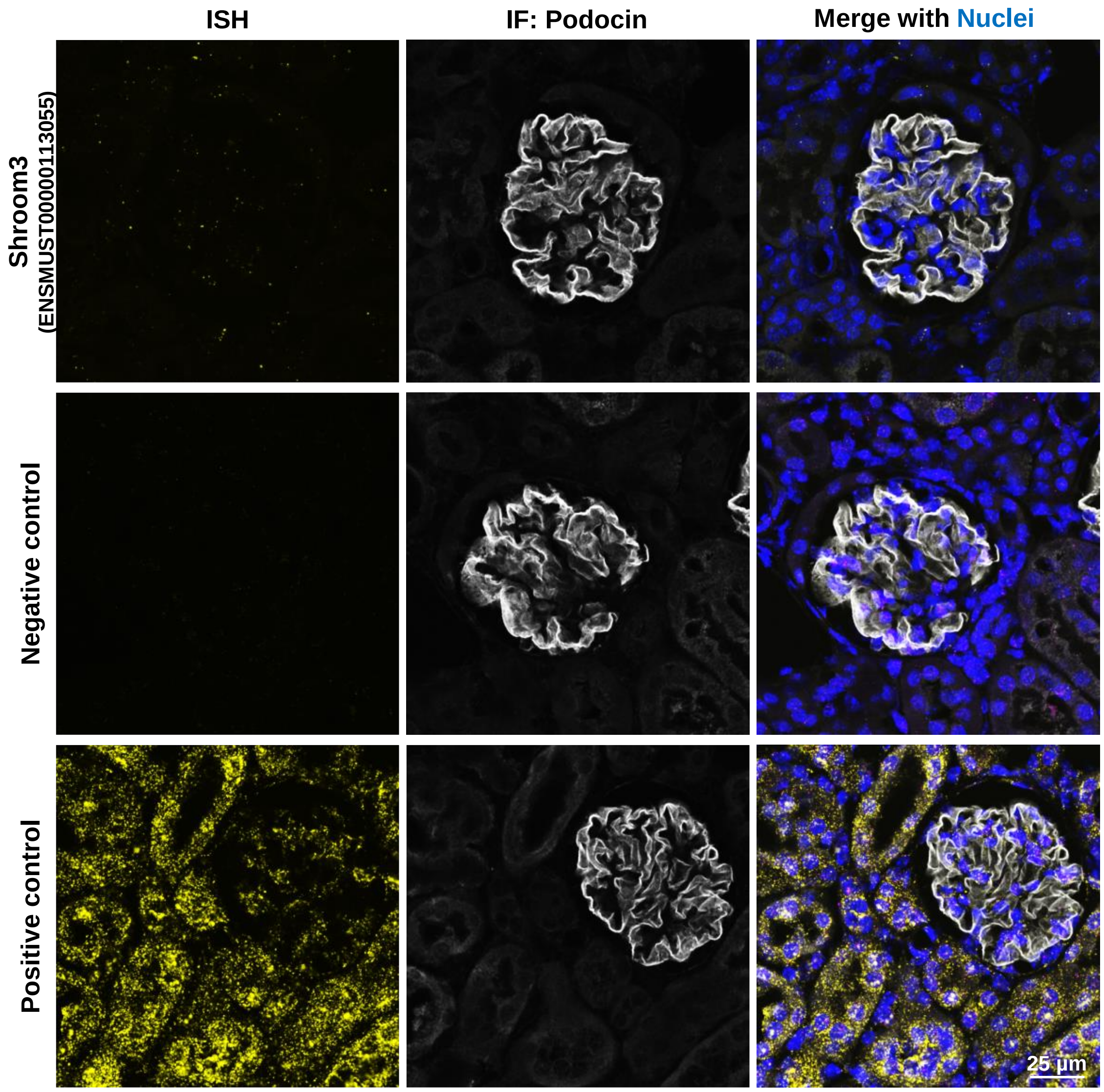

**Fig. S8: RNAscope® ISH Assay.**
*In situ* hybridization of the *Shroom3* isoform ENSMUST00000113055 in mouse kidney (yellow dots). Immunostaining of podocin is shown in white. Nuclei stained with HOECHST (blue). The negative control (*dapB*) and the positive control (*Ppib*) are also shown. Scale bar represents 25 µm.

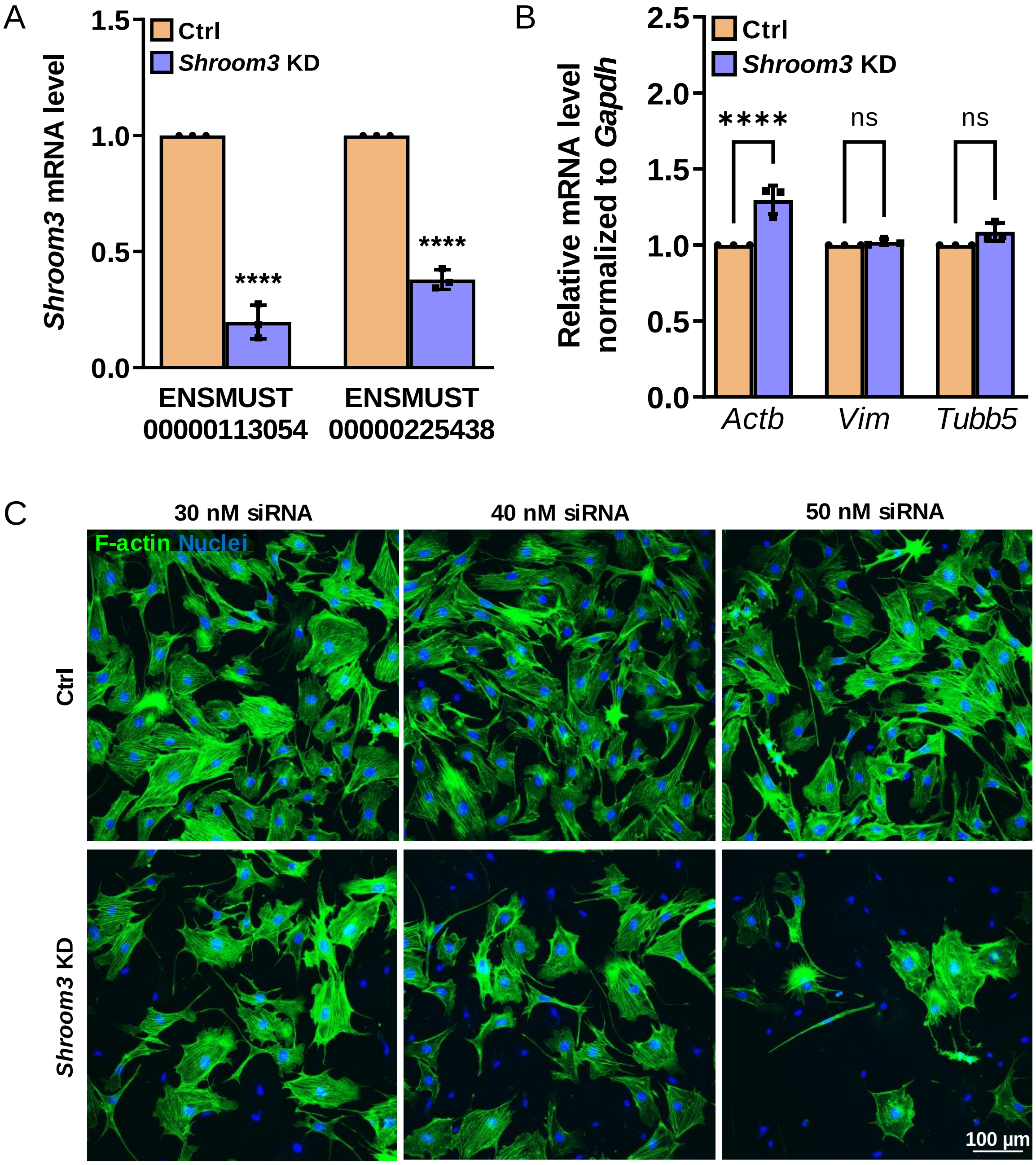

**Fig. S9: *Shroom3* Knockdown (KD) in cultured murine podocytes**

(A) qRT-PCR quantification of *Shroom3* knockdown (*Shroom3* KD) podocytes showed a significant decrease of *Shroom3* coded transcript expression levels compared to the control (Ctrl). (B) Beta-Actin (*Actb*) mRNA level was significantly up-regulated in Shroom3 KD podocytes. In contrast, the mRNA level of vimentin (*Vim*) as a marker of intermediate filaments and β-tubulin (*Tubb5*) were not changed significantly. Data are normalized against Ctrl KD and Gapdh and presented as means ± SD. (C) With an increased siRNA concentration the number of cells without visible F-actin increased dramatically in *Shroom3* KD podocytes. F-Actin is shown in green. Nuclei were stained with DAPI (blue). Scale bar represents 100 µm. ** $p$<0.01; **** $p$<0.0001.

### Supplementary Table 1:

Used (q)RT-PCR Primer

| Detected gene | Sequence / Identity |
|---|---|
| *Gapdh* | F: GTGCTGAGTATGTCGTGGAG<br>R: TGGTGCAGGATGCATTGCTG |
| *Myl6*<br>ENSMUST00000218127 | F: GCGAGAAGATGACAGAGGAAG<br>R: TCACCCCGACAGGATATGCCT |
| *Myl6*<br>ENSMUST00000164181 | F: ATGGGTGCTGAAATCCGTCA<br>R: CCATCCGGACAAGCTCTTCA |
| *Shroom3*<br>ENSMUST00000113054 | F: CAGACGCATCCAGCAGATCAC<br>R: ACCATGTTGCTTTCCGGTGC |
| *Shroom3*<br>ENSMUST00000225438 | F: GCTCTCGTGCTTCCCAGATTT<br>R: CACCCCTCCTGACCATGTTG |
| *Shroom3*<br>ENSMUST00000113055 | F: CTCTTGCCTCCATCTGGGTC<br>R: GGCTTTGCCCCCTTCTTCAA |
| *Tes* | F: GGGTGACGTGAAGTTTCCCT<br>R: AGCCAGCCCTTTCAGCATAG |
| *Serpine1* | F: TGCAAAAGGTCAGGATCGAGG<br>R: GTGCCGAACCACAAAGAGAAAG |
| *Tagln2* | F: TGGCATTAACACCACGGACA<br>R: CAGAGAAGAGCCCATCGTCC |
| *Nes* | F: GAGAGGCGCTGGAACAGAGA<br>R: CTGCCCACCTTCCAGGATCT |
| *Ezr* | F: TATGCCGTTCAGGCCAAGTT<br>R: CTGGTCCCTGCTGAGCTTGT |
| *Nrip3* | F: AGTCACCCTACTGAGGACCG<br>R: ATGGCGTCATGGAACCCTTC |
| *Npr2* | F: GACCCAGCACAACCAGATTTC<br>R: GTTCTACCTGAGTTGGTGGCT |

### Supplementary Table 2:

#### LC-MS/MS parameter (data independent mode; quantitative data)

| ***Data independent analyses (DIA)*** | |
|---|---|
| ***reversed phase liquid chromatography*** | **Ultimate 3000 RSLC (Thermo Scientific)** |
| *Trap column* | 75 µm inner diameter, packed with 3 µm C18 particles (Acclaim PepMap100, Thermo Scientific) |
| *Analytical column* | 75 µm inner diameter, packed with 2.6 µm C18 particles (Accucore, 25 cm, Thermo Scientific) |
| *Flow rate* | 300 nl/min |
| *column oven temperature* | 40°C |
| *buffer system* | binary buffer system consisting of 0.1% acetic acid in HPLC-grade water (buffer A) and 100% ACN in 0.1% acetic acid (buffer B) |
| *gradient* | gradient of buffer B: 2min 2% to 5 %, 8min 7%, 60min 7% to 25%, 5min 25 to 40%, 2 min 40% to 90%, 6 min 90%, 2 min 90% to 2%, 10 min 2% |
| ***Mass spectrometer*** | Exploris 480 |
| *operation mode* | data-independent |
| *electrospray* | Nanospray Flex Ion Source |
| ***Full MS*** | |
| *MS scan resolution* | 120,000 |
| *Normalized AGC target* | 300 % |
| *maximum ion injection time for the MS scan* | 60 ms |
| *Scan range* | 350 to 1200 *m/z* |
| *Spectra data type* | profile |
| ***dd-MS2*** | |
| *Resolution* | 30,000 |
| *Normalized MS/MS AGC target* | 3000 |
| *maximum ion injection time for the MS/MS scans* | auto |
| *Spectra data type* | profile |
| *selection for MS/MS* | 1 |
| *isolation window* | 65 windows *m/z* 13 (overlap *m/z* 2) |
| *Fixed first mass* | 200 |
| *dissociation mode* | higher energy collisional dissociation (HCD) |
| *normalized collision energy* | fixed, 30 |
| *dissociation mode* | HCD |

**Spectronaut parameters for peptide/protein identification and intensity extraction**

| | |
|---|---|
| Spectronaut 17.0.221202.55965 | |
| Computer Name: AGVOE-SPECTRONA | |
| User Domain Name: AGVOE-SPECTRONA | |
| User Name: spectronaut | |
| Analysis Mode: UI | |
| Analysis Type: directDIA | |
| Analysis Date: 17-December-2022 | |
| Database: Mus-musculus_04-2022.fasta (17,107 entries) | |
| Settings Used: C_FunGene_directDIA_sparse_no_imputing | |
| DIA Analysis\Calibration | |
| MZ Extraction Strategy: | Maximum Intensity |
| Allow source specific iRT Calibration: | True |
| Precision iRT: | True |
| Exclude De-amidated Peptides: | True |
| iRT <-> RT Regression Type: | Local (Non-Linear) Regression |
| MS1 Mass Tolerance Strategy: | System Default |
| MS2 Mass Tolerance Strategy: | System Default |
| DIA Analysis\Identification | |
| Precursor Qvalue Cutoff: | 0.001 |
| Precursor PEP Cutoff: | 0.2 |
| Protein Qvalue Cutoff (Experiment): | 0.01 |
| Protein Qvalue Cutoff (Run): | 0.05 |
| Protein PEP Cutoff: | 0.75 |
| Single Hit Definition: | By Stripped Sequence |
| Exclude Single Hit Proteins: | False |
| Exclude Duplicate Assays: | True |
| Exclude Predicted Fragment Scores: | False |
| Generate Decoys: | True |
| Decoy Generation Method: | Mutated |
| Preferred Fragment Source: | NN Predicted Fragments |
| Decoy Limit Strategy: | Dynamic |
| Library Size Fraction: | 0.1 |
| Pvalue Estimator: | Kernel Density Estimator |
| DIA Analysis\Pipeline Mode | |
| Generate SNE File: | True |
| Store Ion traces in SNE: | False |
| Post Analysis Reports: | |
| CV Density Line Chart: | True |
| CVs Below X Bar Chart: | True |
| Data Completeness Bar Chart: | True |
| Run Identifications Bar Chart: | True |
| Scoring Histograms: | True |
| Report Schema: | C_FunGene_complex (Normal) |
| Reporting Unit: | Across Experiment |
| DIA Analysis\Post Analysis | |

| | |
|---|---|
| Differential Abundance Testing: | NA |
| Group-Wise Testing Correction: | False |
| Differential Abundance Grouping: | Major Group (Quantification Settings) |
| Smallest Quantitative Unit: | Precursor Ion (Quantification Settings) |
| Use All MS-Level Quantities: | False |
| Calculate Explained TIC: | None |
| Calculate Sample Correlation Matrix: | True |
| Hierarchical Clustering: | True |
| Distance Metric: | Manhattan Distance |
| Linkage Strategy: | Ward's Method |
| Order Runs by Clustering: | True |
| Z-score Transformation: | False |
| DIA Analysis\Protein Inference | |
| Protein Inference Workflow: | Automatic |
| Inference Algorithm: | IDPicker |
| DIA Analysis\PTM Workflow | |
| PTM Localization: | False |
| DIA Analysis\Quantification | |
| Precursor Filtering: | Identified (Qvalue) |
| Imputation Strategy: | Use Background Signal |
| Proteotypicity Filter: | None |
| Protein LFQ Method: | MaxLFQ |
| Quantity MS Level: | MS2 |
| Quantity Type: | Area |
| Cross-Run Normalization: | True |
| Normalization Filter Type: | None |
| Normalization Strategy: | Local Normalization |
| Row Selection: | Identified in at least 1 Run (Sparse) |
| Interference Correction: | True |
| Only Identified Peptides: | True |
| Exclude All Multi-Channel Interferences: | True |
| MS1 Min: | 2 |
| MS2 Min: | 3 |
| Major (Protein) Grouping: | by Protein Group Id |
| Minor (Peptide) Grouping: | by Stripped Sequence |
| Major Group Quantity: | Mean peptide quantity |
| Major Group Top N: | True |
| Max: | 3 |
| Min: | 2 |
| Minor Group Quantity: | Sum precursor quantity |
| Minor Group Top N: | False |
| DIA Analysis\Workflow | |
| Method Evaluation: | False |
| MS2 DeMultiplexing: | Automatic |

| | |
|---|---|
| Profiling Strategy: | iRT Profiling |
| Carry-over exact Peak Boundaries: | False |
| Profiling Row Selection: | Minimum Qvalue Row Selection |
| Qvalue Threshold: | 0.001 |
| Profiling Target Selection: | Profile only non-identified Precursors |
| Identification Criterion: | Qvalue |
| Threshold: | 0.001 |
| Run Limit for directDIA Library: | -1 |
| Unify Peptide Peaks Strategy: | Select corresponding Peak |
| DIA Analysis\XIC Extraction | |
| XIC IM Extraction Window: | Dynamic |
| Correction Factor: | 1 |
| XIC RT Extraction Window: | Dynamic |
| Correction Factor: | 1 |
| MS1 Mass Tolerance Strategy: | Dynamic |
| Correction Factor: | 1 |
| MS2 Mass Tolerance Strategy: | Dynamic |
| Correction Factor: | 1 |
| Pulsar Search\Identification | |
| PSM FDR: | 0.01 |
| Peptide FDR: | 0.01 |
| Protein Group FDR: | 0.01 |
| PTM Localization Filter: | False |
| Pulsar Search\Modifications | |
| Max Variable Modifications: | 5 |
| Select Modifications: | |
| Fixed Modifications:: | Carbamidomethyl (C) |
| Variable Modifications: : | Acetyl (Protein N-term), Oxidation (M) |
| Pulsar Search\Peptides | |
| Enzymes / Cleavage Rules: | Trypsin/P |
| Digest Type: | Specific |
| Max Peptide Length: | 52 |
| Min Peptide Length: | 7 |
| Missed Cleavages: | 2 |
| Toggle N-terminal M: | True |

### Supplementary methods

#### Cell Culture

Podocytes were maintained in RPMI 1640 medium (Sigma-Aldrich, St. Louis, MO, USA) supplemented with 10% fetal bovine serum (FBS; Boehringer Mannheim, Mannheim, Germany), 100 U/ml penicillin, and 0.1 mg/ml streptomycin (Thermo Fisher Scientific, Waltham, MA, USA). To propagate podocytes, we cultivated cells at 33°C. To induce differentiation, we maintained podocytes at 38°C for at least two weeks before applying mechanical stretch.

#### RNA extraction, cDNA synthesis and qRT-PCR

Cells were processed in Tri-Reagent (Sigma-Aldrich) according to manufacturer's instructions. For cDNA synthesis, 1 µg of the isolated total RNA was transcribed using the QuantiTect Reverse Transcription Kit (Qiagen, Hilden, Germany). The quantitative real-time PCR (qRT-PCR) analysis was performed on a QuantStudio™ 5 Real-Time PCR System (Thermo Fisher Scientific) using the iTaq Universal SYBR Green Supermix (Bio-Rad) with *Gapdh* as reference gene. Relative quantifications of the mRNA levels were done by the efficiency corrected calculation model by Pfaffl and are shown with standard deviations (SD) from at least three biological replicates. Used primers are compiled in Supplemental Table 1.

#### Isolation of murine glomeruli

To analyze individual glomeruli, the kidneys were removed, and chopped up with scissors with 1 ml of cold Hank's Balanced Salt Solution (HBSS; Thermo Fisher Scientific). The cell suspension was then centrifuged at 300 × g for 5 min at 4°C. The supernatant was discarded, the cell pellet was digested with 1 ml of collagenase A (5 mg/ml) (Roche, Basel, Switzerland) and washed with 3 ml cold HBSS at 37°C for 15 min with gentle agitation. The reaction was stopped with 4 ml of RPMI 1640 (Sigma-Aldrich) without phenol supplemented with 10% fetal bovine serum (Thermo Fisher Scientific). The digested tissue was spun down at 300x g for 5 min at 24°C, the supernatant was discarded and the pellet was dissolved in 1 ml of HBSS. The digested tissue was then gently pressed through a 150 µm cell strainer (pluriSelect, Leipzig, Germany) and large tissue particles were pressed through the strainer with the rubber plunger of a 1-ml syringe. The procedure was repeated with cell strainers of different diameters (100 µm and 40 µm). The glomerular and tubular fragments are retained above 40 µm. The supernatant containing highly purified glomeruli was collected and centrifuged at 300x g at 24°C for 5 min. The supernatant was then removed and in 1 ml Tri-Reagent (Sigma-Aldrich) was added.

#### Immunofluorescence staining

Cells were fixed using 2% PFA (paraformaldehyde) in PBS (phosphate buffered saline) for 10 min at room temperature (RT), washed and subjected to blocking solution (PBS, 2% fetal bovine serum, 2% bovine serum fraction V, 0.2% fish gelatine) for 1 hour at RT. Primary antibodies were incubated for 60 min at RT. Actin was stained using Alexa Fluor® phalloidin (Thermo Fisher Scientific). After a washing step with PBS (3 × 3 min) cells were incubated with secondary antibodies for 40 min at RT. Bound antibodies were visualized with Cy2- or Cy3-conjugated secondary antibodies (Jackson ImmunoResearch Laboratories, West Grove, USA). For nuclei staining DAPI (Sigma-Aldrich, 1 µg/ml) was used for 5 min. All samples were mounted in Mowiol (Carl Roth, Karlsruhe, Germany) and used for laser scanning microscopy (LSM). For paraffin sections, samples were dehydrated and embedded in paraffin by standard procedures. Paraffin sections (5 µm) were cut on a Leica SM 2000R (Leica Microsystems, Wetzlar, Germany). After rehydration, sections were unmasked in Tris (pH 9.0) or in citrate buffer (0.1 M, pH 6.0) by heating for 5 min in a pressure cooker. The sections were stained with 1 µg/ml DAPI (Sigma-Aldrich) for 5 min. For immunofluorescence double-staining, samples were incubated with an antibody against MYL6 (PA5-106803; from Thermo Fisher Scientific) and synaptopodin (61094, Progen Biotechnik GmbH, Heidelberg, Germany) overnight. Samples were washed with 1x PBS for 3x5 min and incubated with Cy2- and Cy3-conjugated secondary antibodies (Jackson ImmunoResearch Laboratories) for 1 hour. After additional washing, the samples were mounted in Mowiol (Carl Roth) for fluorescence microscopy.

#### Histology on human kidney biopsies

Kidney biopsies were received from the Department of Nephropathology, Institute of Pathology, University Hospital Erlangen, Germany. The use of remnant kidney biopsy material was approved by the Ethics Committee of the Friedrich-Alexander-University of Erlangen-Nuremberg, waiving the need for retrospective consent for the use of archived rest material (Re.-No.4415 and Re.-No.22-150-D). Sample fixation and preparation of the control, HN, DN, and FSGS group were identical.

#### Combined multiplex fluorescence *in situ* hybridization (RNAscope) and immunofluorescence

RNA *in situ* hybridization was performed for Shroom3 isoforms using RNAscope Multiplex Fluorescent V2 kit (Advanced Cell Diagnostics, Inc.) with the following modifications. Tissue sections were deparaffinized in xylene (two times for 5 min) and rehydrated in 100% ethanol (two times for one minute). The pre-treatment procedure included the application of RNAscope Hydrogen Peroxide (10 min at room temperature), the submersion in boiling RNAscope Target Retrieval reagent (15 min), and the treatment with RNAscope Protease Plus (15 min at 40°). RNA scope complementary probes were hybridized for 2 hours at 40°C in the ACDbio HybEZ oven. Each probe has been assigned a different channel: Shroom3_054 (Cat No. 1276721-C1), Shroom3_055 (Cat No. 1077361-C2) and Shroom3_438 (Cat No. 1298351-C3). 3-plex Positive (Polr2a, Ppib and Ubc) and 3-plex Negative Control (dapB) probes were used as positive control and negative control, respectively. Sections were stored in 5× SSC (Invitrogen) overnight. The hybridization signals were then amplified and detected using Opal 520, 570 and 690 dyes diluted 1:1500 in TSA amplification buffer (ACDbio). After the detection steps, sections were subjected to blocking solution (PBS, 2% fetal bovine serum, 2% bovine serum fraction V, 0.2% fish gelatine) for 1 hour at RT (room temperature) and incubated with primary antibody against podocin (1:150) (IBL International GmbH, Hamburg, Germany). Primary NPHS2 antibodies were detected using AlexaFluor 750 secondary antibodies. Slides were counterstained with DAPI (Sigma-Aldrich, 1 µg/ml) and mounted in Mowiol (Carl Roth) and imaged by confocal laser scanning microscopy.

##### Laser scanning microscopy

Images were captured with an Olympus FV3000 confocal microscope (Olympus, Tokyo, Japan) with 20x/40x/60x oil immersion objectives and Olympus FV3000 CellSense software.
